# A Whole-Mouse-Brain Model of Coupled Neural Masses: Analysis of Network Dynamics

**DOI:** 10.64898/2026.10.29.762816

**Authors:** Christopher J Parker, Nicholas M Timme

**Affiliations:** Department of Biomedical Sciences, University of Cincinnati, 2600 Clifton Ave, Cincinnati, 45221, Ohio, USA

**Keywords:** Neural mass model, Jansen-Rit model, mouse brain, dynamical systems analysis

## Abstract

Neural mass models coupled through empirical structural connectomes are widely used to simulate whole-brain activity, but the dynamics of the resulting large, delay-coupled networks are rarely analyzed directly. We present a dynamical systems analysis of 434 Jansen-Rit neural masses coupled by the Allen Mouse Brain Connectivity Atlas at the canonical single-node parameter values. Pseudo-arclength continuation of the network equilibria in the global coupling scale and in the baseline external input, with stability determined from the characteristic roots of the delayed linearization by Beyn’s contour-integral method, indicates where the network must oscillate and where it can rest. Simulations show that the oscillatory regime is chaotic: the maximum Lyapunov exponent is positive for the empirical connectome, grows with network size and coupling scale, and is insensitive to the conduction delays, but it is suppressed when the arrangement of connection weights is flattened, shuffled, or rewired. The same weight modifications that suppress chaos drive the network toward full synchrony and abolish metastability, as measured by the Kuramoto order parameter. We discuss the consequences of these findings for fitting whole-brain neural mass models to empirical data.

## 1 Introduction

Models of vastly different scales are routinely used in computational neuroscience: from extremely detailed models of individual neurons (e.g. Hay, Hill, Schürmann, Markram, & Segev, 2011), to networks of comparatively simplified neurons (e.g. Izhikevich & Edelman, 2008), up to various types of neural mass models (e.g. Jansen & Rit, 1995; Wilson & Cowan, 1972; Wong & Wang, 2006). Neural mass models are an abstract representation of brain dynamics, where the state variables represent aggregate measures of activity such as average postsynaptic potential (PSP). These models have been developed to match brain-wide dynamics such as EEG or BOLD signal, which represent activity averaged across many neurons. They are attractive to neuroscientists because they are significantly less computationally intensive when compared with large networks of individual neurons, their dynamics are easier to study, and they can be related to non-invasive neural data gathered in humans. It remains infeasible to simulate billions of neurons concurrently to represent a full brain, while a network of coupled neural masses can reliably reproduce certain characteristics of brain-wide activity. Importantly, a model that can reproduce empirical data gives us a hypothesis (the model parameters) about the conditions necessary for the observed data to arise.

One such neural mass model is that by Jansen and Rit (1995), which utilizes three interacting sub-populations of neurons to represent an abstract cortical column: excitatory pyramidal neurons, excitatory interneurons, and inhibitory interneurons. The pyramidal neurons drive both interneuron populations, and also drive other coupled columns. The interneuron populations only connect within a single column, and only to the pyramidal neurons, creating positive (excitatory) and negative (inhibitory) feedback loops. The outputs are PSPs in mV, originally designed to model the EEG alpha rhythm (around 10 Hz). Dynamics of the Jansen-Rit model have been well-studied in the literature for single-node and few-node cases (Ahmadizadeh et al., 2018; Grimbert & Faugeras, 2006; Huang, Zhang, Meng, & Zhu, 2011; Jansen & Rit, 1995; Spiegler, Kiebel, Atay, & Knösche, 2010; Touboul, Wendling, Chauvel, & Faugeras, 2011). Modeling the many-node case is becoming more feasible as available computing resources scale up and structural connectome data becomes more readily available, but the dynamical systems analysis literature remains limited in this case (Forrester, Crofts, Sotiropoulos, Coombes, & O’Dea, 2020).

Here, we present dynamical systems analyses of a network of coupled Jansen-Rit neural masses (Jansen & Rit, 1995) connected with Allen Institute Mouse Brain Connectivity Atlas data (Allen Institute for Brain Science, 2017; Knox et al., 2018; Oh et al., 2014). In all tests, we have used the canonical parameter values for the individual nodes, and we have based our default connectivity-related parameters on those of The Virtual Brain and The Virtual Mouse Brain (Melozzi, Woodman, Jirsa, & Bernard, 2017; Sanz Leon et al., 2013). The model consists of 217 regions per hemisphere (434 total neural masses), with the network mirrored across hemispheres.

Bifurcation analysis is a key tool when attempting to understand a dynamical system like the Jansen-Rit model. In general, bifurcations are points where the behavior of the system suddenly changes as we smoothly vary parameter(s) of interest. Bifurcation analysis is useful because it can help us determine all possible behaviors of the system, as opposed to typical simulations which only reveal which behaviors the system happened to exhibit in a given run. In the single-node case, as presented by Grimbert and Faugeras (2006), the model behavior is organized around a few important points when varying the external input to the node. Of primary interest in the analyses we report here: the system exhibits bistability for low external input (low- or high-activity stable steady states, between a pair of fold bifurcations), limit cycle oscillations between two Hopf bifurcations for moderate external input (the limit cycle gives roughly 10 Hz oscillations), and a single stable steady state for high external input.

We present a network bifurcation analysis of pyramidal neuron postsynaptic potential (PSP) on the global coupling scale parameter (*G*) and the baseline external input to each node (*µ*). This can be viewed as an extension of the analysis in Grimbert and Faugeras (2006) to the many-node case, where *µ* and *G* are both related to the bifurcation parameter of their analysis: total external input *p* (*µ* represents external input from unmodeled structures, *G* scales the external input from other modeled nodes). Given that the network structure turns the ordinary differential equation (ODE) system into a delay differential equation (DDE) system, bifurcation analysis becomes significantly more complex and computationally expensive. We use pseudo-arclength continuation (PALC) to find equilibrium curves. Eigenvalues of the system along these curves are tracked to assess stability of the system and detect bifurcations (due to the nature of DDEs, eigenvalues are tracked only in a box of interest, see Section 2 for more information). This allows us to place reasonable bounds on *G* and *µ* where we expect the system to behave in a desired way (i.e. oscillating with moderate synchrony between nodes).

Chaos (sensitive dependence on initial conditions) is another important property of some dynamical systems. In a chaotic system, two solutions started from infinitesimally-close points will diverge exponentially over time, and therefore the system behaves unpredictably over long horizons. Given the results of Huang et al. (2011) showing chaotic dynamics for specific parameter regimes in the case of two coupled nodes, we present a grid search over combinations of global coupling strength *G*, number of networked nodes, and several types of connectivity modification. Our aim was to determine if the default, alpha-rhythm operating regime exhibits chaotic dynamics for large networks of nodes. On finding that this is the case for the standard parcellation of the Allen Institute mouse connectome, we wondered whether that chaos is dependent on specific distributions of weights and delays in the network, or if generic networks of coupled nodes without physiological basis also exhibit the same behavior. To assess chaotic dynamics in our model, we compute estimates of the maximum Lyapunov exponent (MLE) via the single-trajectory method of Farmer (1982). We found that the networked system exhibits chaotic dynamics at the canonical parameter values of Jansen and Rit (1995) without any noise in the system. Given the difficulty chaotic dynamics introduce into numerical optimization procedures for fitting the model system to empirical data, we also present some practical suggestions for working with models of this type.

Finally, we present the results of a grid search procedure on global coupling scale and axonal conduction velocity that examines synchrony and metastability in the network, the definitions of which follow Deco, Kringelbach, Jirsa, and Ritter (2017) in our work (mean and standard deviation of Kuramoto order parameter *R*(*t*)). The presence of metastability in the brain has been suggested across several data modalities including fMRI (e.g., Alderson, Bokde, Kelso, Maguire, & Coyle, 2020; Alderson et al., 2018), MEG (e.g., Baker et al., 2014; Cabral et al., 2022), and intracortical recordings (e.g., La Camera, Fontanini, & Mazzucato, 2019; Mazzucato, Fontanini, & La Camera, 2015). See (Brinkman et al., 2022; Hancock et al., 2025) for recent reviews on metastability. Interestingly, we found that network modifications that decrease the MLE also tend to decrease metastability while increasing synchrony.

Overall, our analyses of this whole-mouse-brain neural mass model provide insights into the behavior of the network at various levels of external input, the cause of chaotic dynamics exhibited in the default operating regime, and the relation between the chaotic dynamics and network synchrony and metastability. These results will directly impact future whole-brain modeling studies and extensions of these approaches to human-derived models and data.

## 2 Methods

Here we present descriptions of the mathematical techniques used in this research. For additional information, we direct the reader to the code archive (publicly available at https://doi.org/10.60696/uc.34845648), where there are verbose comments alongside the code.

### 2.1 Jansen-Rit Model Description

The Jansen-Rit neural mass model (Jansen & Rit, 1995) consists of three coupled subpopulations of neurons: excitatory pyramidal neurons (*y*_0_ represents EPSP from pyramidal neurons to both interneuron populations), excitatory interneurons (*y*_1_ represents EPSP from excitatory interneurons to pyramidal neurons), and inhibitory interneurons (*y*_2_ represents IPSP from inhibitory interneurons to pyramidal neurons). For the full Jansen-Rit coupled network, node *j* ∈ [1, *N* ] obeys

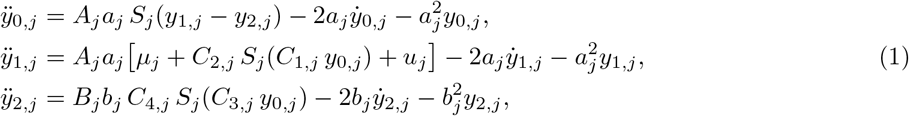

with sigmoid

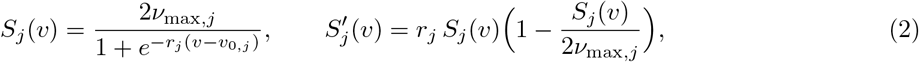

and network input

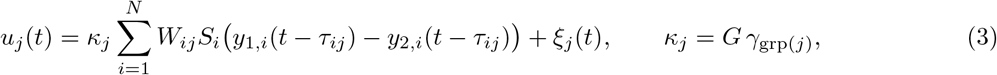

for target node *j*; coupling scale *κ*_*j*_ (which is a scalar *G* that does not vary by node in the current tests, although the code allows for a gain *γ*_grp(*j*)_ on the functional group grp(*j*) to which node *j* belongs); connectivity weight matrix *W* with *W*_*ij*_ representing the connectivity strength from node *i* to node *j*; delays *τ* where *τ*_*ij*_ = *D*_*ij*_*/v* represents the axonal conduction delay from node *i* to node *j* for inter-centroid distance *D*_*ij*_ and conduction speed *v*; and *ξ*_*j*_(*t*) an external stimulation or random noise (always zero in the current tests). See Table 1 for the parameters in the model, their default values (or ranges tested in this work), and a brief description of the physiological meaning of each.

**Table 1.**
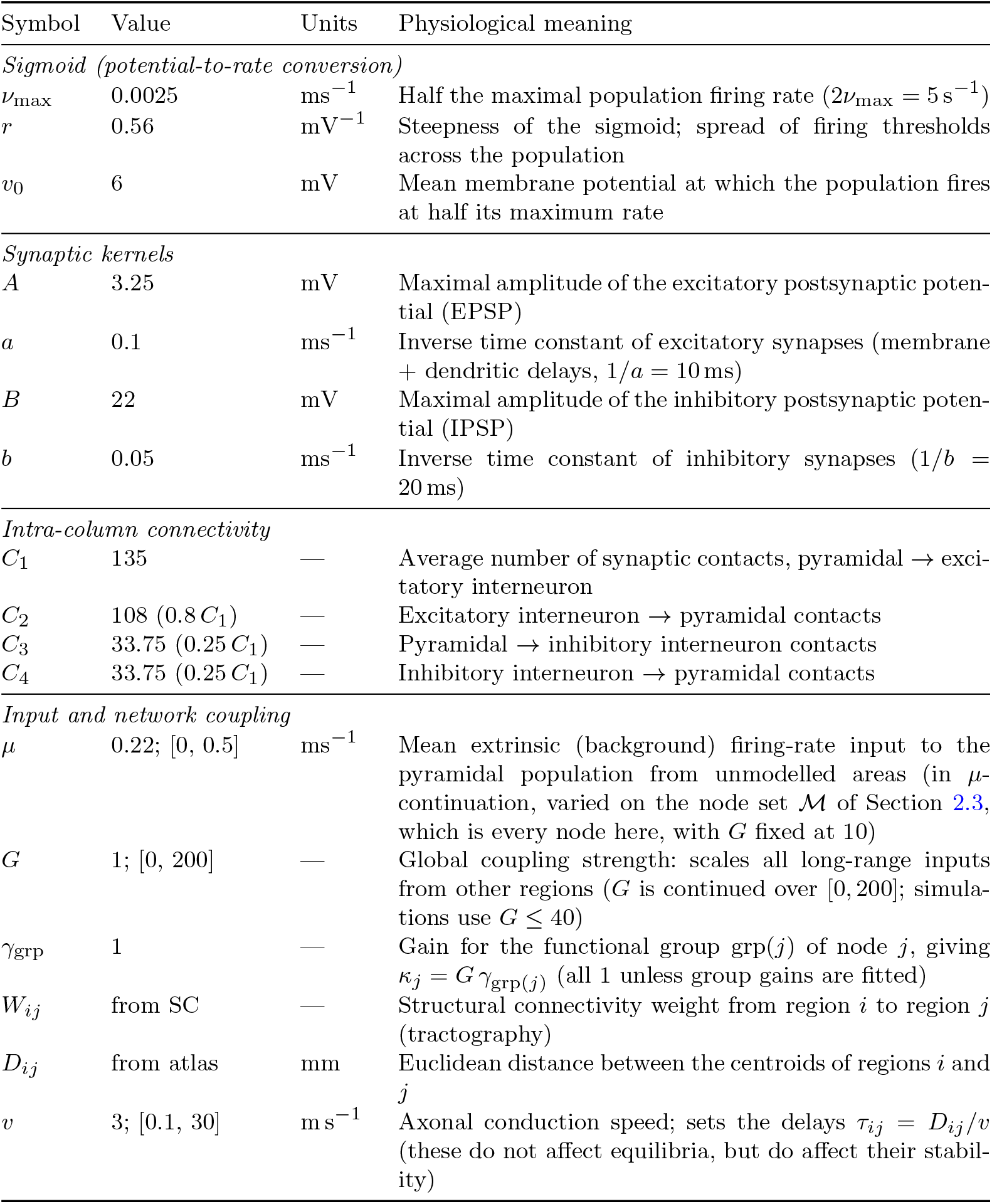
Jansen–Rit network model parameters. Local parameters are identical across all *N* = 434 regions and follow Jansen & Rit (1995), except *µ*, which follows The Virtual Brain. Time is in ms throughout. All parameters have their default value listed; we additionally list ranges for *µ, G*, and *v* that cover the values tested in analyses.

| Symbol | Value | Units | Physiological meaning |
|---|---|---|---|
| *Sigmoid (potential-to-rate conversion)* |  |  |  |
| *ν*<sub>max</sub> | 0.0025 | ms<sup>−1</sup> | Half the maximal population firing rate (2*ν*<sub>max</sub> = 5 s<sup>−1</sup>) |
| *r* | 0.56 | mV<sup>−1</sup> | Steepness of the sigmoid; spread of firing thresholds across the population |
| *v*<sub>0</sub> | 6 | mV | Mean membrane potential at which the population fires at half its maximum rate |
| *Synaptic kernels* |  |  |  |
| *A* | 3.25 | mV | Maximal amplitude of the excitatory postsynaptic poten-tial (EPSP) |
| *a* | 0.1 | ms<sup>−1</sup> | Inverse time constant of excitatory synapses (membrane + dendritic delays, 1/*a* = 10 ms) |
| *B* | 22 | mV | Maximal amplitude of the inhibitory postsynaptic poten-tial (IPSP) |
| *b* | 0.05 | ms<sup>−1</sup> | Inverse time constant of inhibitory synapses (1/*b* = 20 ms) |
| *Intra-column connectivity* |  |  |  |
| *C*<sub>1</sub> | 135 | — | Average number of synaptic contacts, pyramidal → exci-tatory interneuron |
| *C*<sub>2</sub> | 108 (0.8 *C*<sub>1</sub>) | — | Excitatory interneuron → pyramidal contacts |
| *C*<sub>3</sub> | 33.75 (0.25 *C*<sub>1</sub>) | — | Pyramidal → inhibitory interneuron contacts |
| *C*<sub>4</sub> | 33.75 (0.25 *C*<sub>1</sub>) | — | Inhibitory interneuron → pyramidal contacts |
| *Input and network coupling* |  |  |  |
| *μ* | 0.22; [0, 0.5] | ms<sup>−1</sup> | Mean extrinsic (background) firing-rate input to the pyramidal population from unmodelled areas (in *μ*-continuation, varied on the node set ℳ of Section 2.3, which is every node here, with *G* fixed at 10) |
| *G* | 1; [0, 200] | — | Global coupling strength: scales all long-range inputs from other regions (*G* is continued over [0, 200]; simula-tions use *G* ≤ 40) |
| *γ*<sub>grp</sub> | 1 | — | Gain for the functional group grp(*j*) of node *j*, giving *κ*<sub>*j*</sub> = *G* *γ*<sub>grp(*j*)</sub> (all 1 unless group gains are fitted) |
| *W*<sub>*ij*</sub> | from SC | — | Structural connectivity weight from region *i* to region *j* (tractography) |
| *D*<sub>*ij*</sub> | from atlas | mm | Euclidean distance between the centroids of regions *i* and *j* |
| *v* | 3; [0.1, 30] | m s<sup>−1</sup> | Axonal conduction speed; sets the delays *τ*<sub>*ij*</sub> = *D*<sub>*ij*</sub>/*v* (these do not affect equilibria, but do affect their stabil-ity) |

All simulations integrate Equation 1 with Heun’s method (the explicit trapezoidal rule) at a fixed step of Δ*t* = 0.5 ms. The delayed firing rates *S*_*i*_(*y*_1,*i*_(*t* −*τ*_*ij*_) −*y*_2,*i*_(*t* −*τ*_*ij*_)) are read from a ring buffer of past emitted rates; because the delays are not integer multiples of Δ*t*, each delayed rate is linearly interpolated between the two buffered steps that bracket *t* −*τ*_*ij*_. The initial history is constant: every buffered entry is set to the emitted rate of the initial state. The code is described in Section 2.8.

When discussing interactions between subpopulations within a single node, we use the term node-local interactions. Interactions from other nodes will be called network interactions.

The value *y*_1_− *y*_2_ represents the average membrane potential in pyramidal neurons, which is taken as a proxy for EEG signal, and often treated as the primary output of the model system. The related value *S*(*y*_1_ − *y*_2_) represents the average firing rate of pyramidal neurons in the population.

#### First-order form and node-local linearization

The stability and Lyapunov computations below use the first-order form of Equation 1. Let

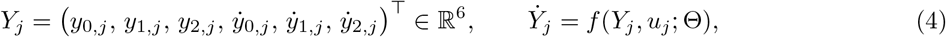

where Θ collects the node parameters of Table 1, so that the full network state is *Y* = (*Y*_1_, …, *Y*_*N*_ ) ∈ ℝ^6*N*^ . At a state *Y*_*j*_ define the sigmoid slopes

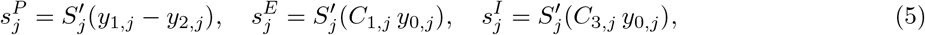

the derivatives of the node-local input firing rates to the pyramidal, excitatory interneuron, and inhibitory interneuron subpopulations, respectively. Linearizing *f* at *Y*_*j*_ with the network input *u*_*j*_ held fixed gives the node-local Jacobian *J*_*j*_, the input sensitivity *c*_*j*_, and the emitted-rate gradient *d*_*j*_:

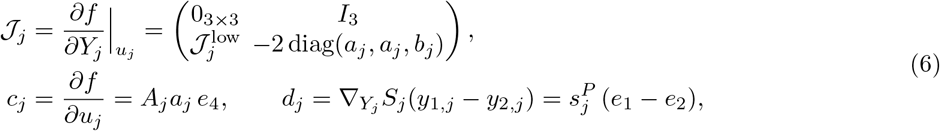

with

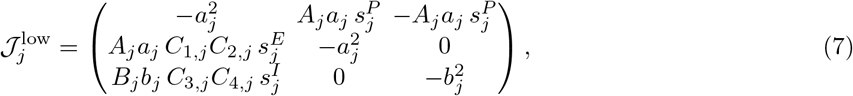

where *e*_0_, …, *e*_5_ is the standard basis of ℝ^6^, indexed from zero to match the ordering of *Y*_*j*_ (the network input enters only the 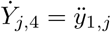 equation). The equilibrium analyses of Section 2.3 utilize the 3*N* unknowns 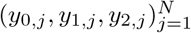 because all derivatives vanish at equilibrium; the stability and Lyapunov analyses of Sections 2.4 and 2.5 utilize the 6*N* -dimensional linearization because stability depends on the full dynamics.

### 2.2 Allen Institute Mouse Brain Connectivity Atlas Description

The Allen Institute Mouse Brain Connectivity Atlas contains data from many axonal tracing experiments in mice (Allen Institute for Brain Science, 2017; Knox et al., 2018; Oh et al., 2014). The primary tracer was an adeno-associated virus (AAV) expressing enhanced green fluorescent protein (EGFP). Precise injections of AAV were performed in targeted locations across many mouse brains, although the data is primarily from injections into the right hemisphere. Imaging was performed using serial two-photon tomography, with the EGFP labeling clearly the locations of axonal projections and their relative strength (through analysis of fluorescence density and intensity across voxels).

For our model, we have used the atlas at its 100 *µ*m isotropic voxel resolution, with all lengths converted to mm. The parcellation we use is the “Brain — Summary Structures” structure tree (ID 167587189) in the Allen SDK (allensdk) Python package; regions in this set without any injection experiment were discarded, leaving *N/*2 = 217 regions per hemisphere. The weight *W*_*ij*_ is the mean projection density over all experiments injected into region *i*, measured in the voxels of region *j* (excluding the injection site itself), so that rows index the injection (source) region and columns index the downstream (target) region, consistent with Equation 3. We have used all injection experiments into each region, regardless of the hemisphere injected; each experiment’s injection hemisphere is determined from its injection coordinate relative to the midline, and the projection density in each target region is classified as ipsilateral or contralateral to that injection. For each source region, the mean ipsilateral density over its experiments gives the within-hemisphere block of *W* and the mean contralateral density gives the between-hemisphere block. The left hemisphere is then taken as the mirror image of the right, so *W* is symmetric under exchange of the hemispheres. This construction is primarily due to the nature of the dataset, but it does have the benefit of making certain analyses more streamlined. The distances *D*_*ij*_ used to set the delays are Euclidean distances between region centroids in the common coordinate framework (mm), with the left-hemisphere centroids mirrored across the midline.

Given that the Jansen-Rit model handles node-local interactions, we have zeroed the diagonal in the weight matrix so that self-connections are given no weight in the network coupling computations. One further pair of entries is zeroed when the connectome is loaded: the nucleus raphe obscurus is a midline structure whose right-hemisphere voxel centroid lies exactly on the midline, so its mirrored left-hemisphere copy has the same centroid, the distance between the two copies is zero, and an edge between them would be a self-connection with no delay. All other region pairs have a positive distance. After these steps, *W* has 177,018 non-zero entries out of 434 ×433 = 187,922 ordered pairs (94%).

In Table 2, we provide descriptive statistics for the connectivity matrix used. In particular, we highlight that the weights are heavily right-skewed (the median is much smaller than the mean), and there is a wide range of in-degree and out-degree values. We note that there are no regions with strong incoming connections from all other regions (connection weight *>* 1× 10^*−*5^), but there are 14 regions (7 regions paired in both hemispheres) that have strong outgoing connections to all other regions. These are (in decreasing order of out-strength): dorsal nucleus raphe (5.143), interpeduncular nucleus (4.491), lateral hypothalamic area (4.167), ventral tegmental area (4.009), laterodorsal tegmental nucleus (2.639), parabrachial nucleus (1.968), and lateral mammillary nucleus (1.407). Note that *W* is mirror-symmetric across hemispheres but not transpose-symmetric: *W*_*ij*_ and *W*_*ji*_ differ in general, because they come from injections into different source regions.

**Table 2.**
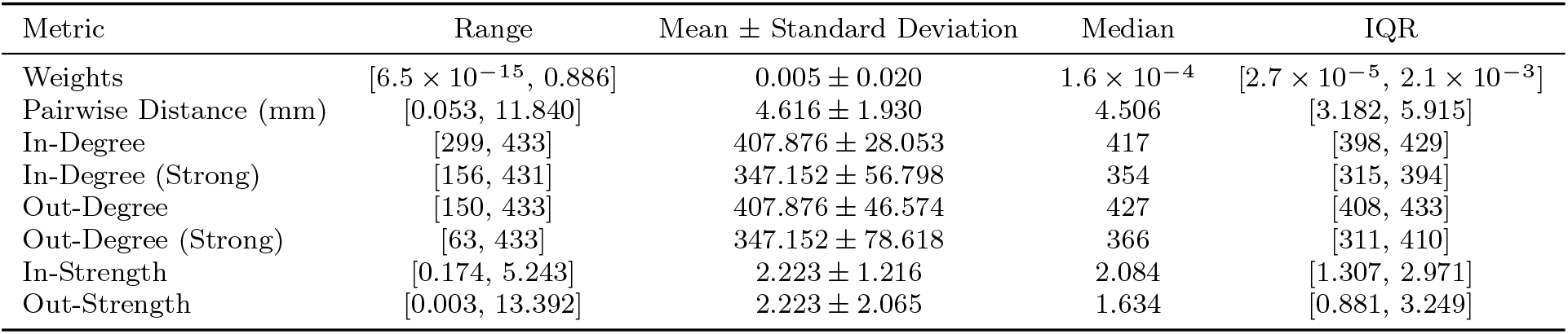
Descriptive statistics for the default connectivity matrix, non-zero entries only (weights and pairwise distances are over the 177,018 non-zero edges). In-Degree and Out-Degree represent the number of non-zero incoming/outgoing connections, respectively. Many regions have in-degree or out-degree of 433, which represents non-zero connection weights to all other regions (including connections with strengths down to 6.5 ×10^*−*15^, which have very little effect). Therefore, we also report In-Degree (Strong) and Out-Degree (Strong) to represent the values when we drop all edges with weights below 1× 10^*−*5^. In-Strength and Out-Strength represent the sum of incoming and outgoing weights. As all edges are counted once as outgoing and once as incoming, the mean values agree between in-/out-degree and in-/out-strength. Mean *±*SD uses the sample standard deviation.

### 2.3 Network Equilibria via PALC

Using pseudo-arclength continuation (PALC), we have traced the network equilibria, reported through the pyramidal firing rate *S*(*y*_1_− *y*_2_) of each node, against global coupling scale *G* and against baseline external input *µ*. See Algorithm 1 for PALC algorithm pseudocode; here we present the equations for the system at equilibrium, the residual equations that we seek roots for in the PALC procedure, and the linearization of the residual system. An important simplification can be made at equilibrium: the derivatives vanish, so the 6*N* first-order differential equations of Equation 4 reduce to 3*N* algebraic equations (three per node) in the unknowns (*y*_0,*j*_, *y*_1,*j*_, *y*_2,*j*_).

#### Jansen-Rit Network at Equilibrium

At an equilibrium all time derivatives vanish, 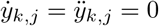 for *k* = 0, 1, 2, *j* ∈ [1, *N* ]. Delays drop out, since *y*_*i*_(*t* − *τ*_*ij*_) = *y*_*i*_ for a constant solution. Then each node satisfies

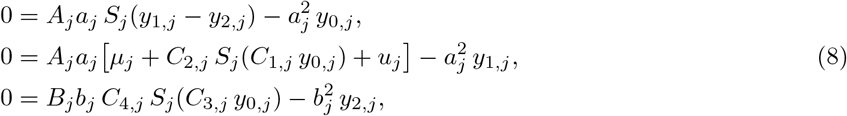

with *u*_*j*_ = *κ*_*j*_ *Σ* _*i*_ *W*_*ij*_ *S*_*i*_(*y*_1,*i*_ − *y*_2,*i*_). Since *a*_*j*_, *b*_*j*_ *>* 0, dividing by 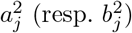 gives the fixed-point form

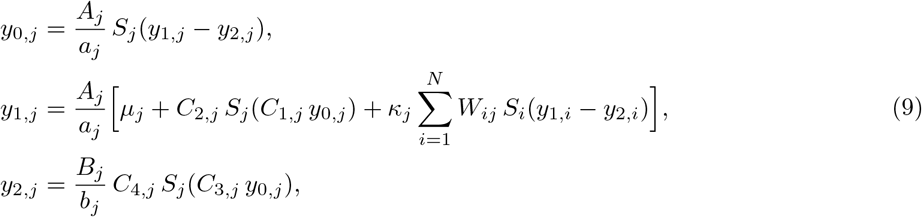

i.e. each equilibrium PSP equals the steady-state gain of its synaptic kernel (*A*_*j*_*/a*_*j*_ or *B*_*j*_*/b*_*j*_) times its input firing rate.

Writing 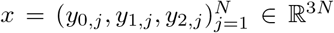 (node-major, so the three unknowns of node *j* are contiguous) and letting *α* ∈ {*G, µ*} be the continuation parameter, we define equilibria as the zeros of the residual *F* : ℝ^3*N*^ × ℝ → ℝ^3*N*^, *F* (*x, α*) = 0, with

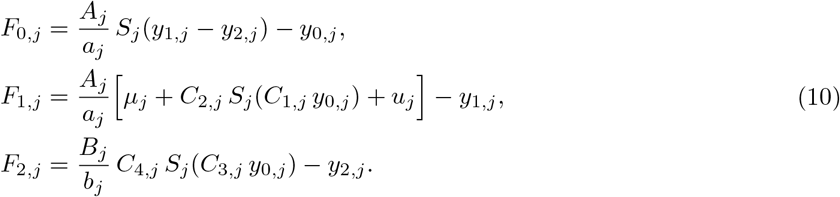

#### State Jacobian F_x_

With the sigmoid slopes 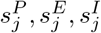 of Equation 5, entries of the Jacobian of the residual system with respect to the state *x* (*F*_*x,j*_ for node *j*) are as follows:

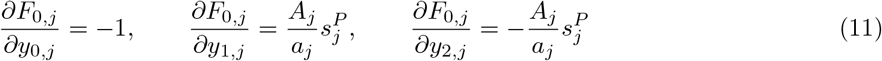

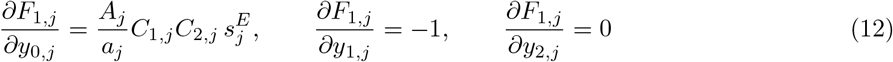

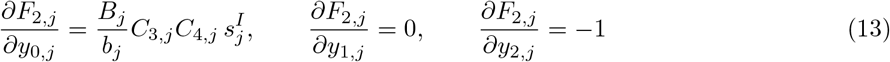

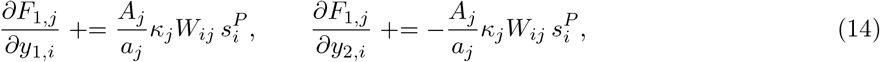

where the *∂F*_1,*j*_*/∂y*_1,*i*_ and *∂F*_1,*j*_*/∂y*_2,*i*_ terms in Equation 14 are accumulated for all *i*∈ [1, *N* ] (although the self-connection term *i* = *j* is zero in all tests here).

In block form, we write the full network Jacobian *F*_*x*_ as

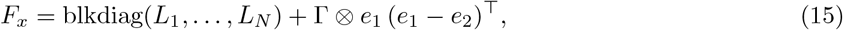

where *L*_*j*_ represents the node-local component of the Jacobian for node *j* (Equations 11, 12, 13 above):

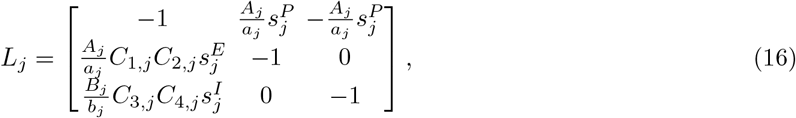

Γ is the coupling term from the rest of the network:

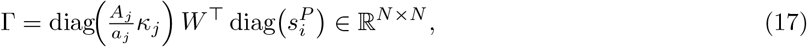

and *e*_0_, *e*_1_, *e*_2_ are the standard basis of ℝ^3^.

#### *Parameter Derivative F*_*α*_

Only *F*_1,*j*_ depends on either *G* or *µ*, so *F*_*α*_ = Σ _*j*_(*∂*_*α*_*F*_1,*j*_) *e*_3(*j−*1)+1_, where *e*_0_, …, *e*_3*N−*1_ is the standard basis of ℝ^3*N*^ indexed from zero to match the node-major ordering of *x*. Depending on whether we are performing continuation in *G* or *µ*, we get different expressions for *F*_*α*_.

Continuation in global coupling, *α* = *G*, gives

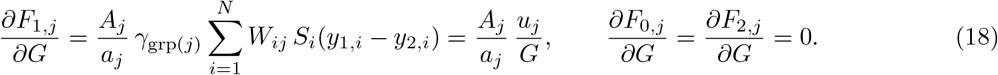

Here, we note again that *κ*_*j*_ = *Gγ*_grp(*j*)_ and *γ*_grp(*j*)_ ≡ 1 in all tests we have run thus far.

Continuation in baseline external input, *α* = *µ*, holds *G* fixed and varies *µ*_*j*_ only on a chosen node set ℳ ⊆ {1, …, *N* } (all nodes in the tests reported here), giving

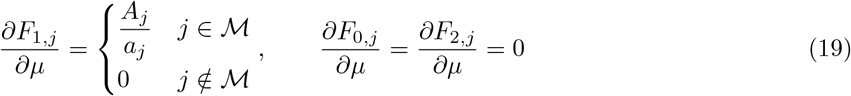

#### PALC Algorithm Description

PALC is differentiated from standard arclength continuation by its ability to follow curves around fold points where the Jacobian *F*_*x*_ becomes singular and following the tangent normally would fail. The PALC algorithm begins by solving the problem

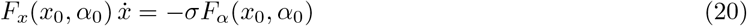

to find the direction 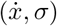 along the equilibrium curve (the null vector of [*F*_*x*_ *F*_*α*_]), which we normalize to obtain our tangent 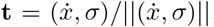. Here, *σ* ∈ {−1, +1} represents the direction in which we step *α* to follow a curve, and we call *z*_0_ = (*x*_0_, *α*_0_) our initial equilibrium state.

Once we have our initial tangent, we take a small step (Δ*s*) in that direction, 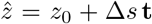. This new point 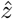 is almost certainly no longer on the curve (because we have taken a step in the best direction for the local, linearized system only), however it is a good guess. We then construct the PALC bordered matrix:

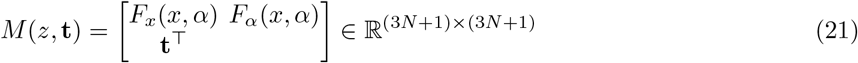

which combines the equilibrium constraint *F* = 0 and the PALC constraint **t**^*⊤*^(*z* −*z*_0_) = Δ*s* (*z* must lie on the hyperplane perpendicular to the tangent through the predictor) through solving the bordered residual equation 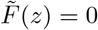 where

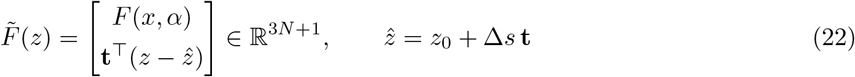

This solution is computed via Newton’s method by solving

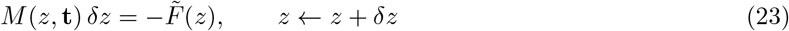

where the addition of the PALC constraint ensures that *M* (*z*, **t**) stays non-singular and we can find a locally unique solution.

The procedure then continues by finding the tangent as in Equation 20, taking a step and correcting back to the curve. We additionally check at each iteration for conditions that indicate points of interest:

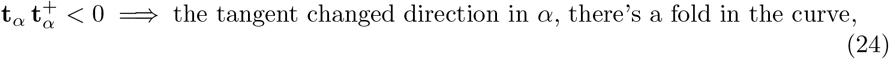

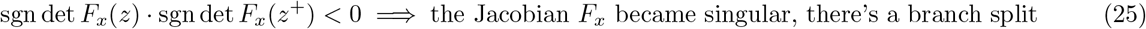

where sgn represents the sign function, **t**_*α*_ is the *α*-component of the unit tangent **t**, and (*z*, **t**) → (*z*^+^, **t**^+^) is the updated curve point after an iteration. Since *F*_*x*_ is also singular at a fold, the second test is only applied when we didn’t change direction in *α* (Algorithm 1). When either test passes, the special point is located by bisection in arclength between *z* and *z*^+^. At a suspected branch point, the null vector of *F*_*x*_ (its right singular vector of smallest singular value) is recorded so that the bifurcating branch can be followed; no branch points were encountered in the runs reported here.

Three implementation details matter for reproducing the curves. First, the parameter coordinate of *z* is scaled by a weight *θ*, i.e. the continuation works with *z* = (*x, θα*) and the bordered matrix carries *F*_*α*_*/θ*. For *G* we use *θ* = 1; for *µ*, to which the equilibrium is far more sensitive (*∂F*_1,*j*_*/∂µ* = *A*_*j*_*/a*_*j*_ = 32.5), *θ* is set to ∥*∂x/∂µ*∥ at the starting point so that an arclength step moves a comparable distance in state and parameter. Second, because *W* and *D* are mirror-symmetric across hemispheres (Section 2.2), the equilibrium we follow is mirror-symmetric, and after every Newton update the state is projected onto the mirror-symmetric subspace, 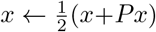 for the hemisphere-exchange permutation *P* ; this removes rounding-error asymmetry that would otherwise accumulate. Third, the runs reported here use an initial step Δ*s* = 0.5, Δ*s*_max_ = 2, Δ*s*_min_ = 10^*−*9^, Newton tolerance *ε*_N_ = 10^*−*10^ on ∥*F* ∥_*∞*_ with at most *k*_max_ = 8 iterations per correction, and the turn threshold *c*_min_ = 0.95 of Algorithm 1. The starting equilibrium at *G* = 0 (or at the starting *µ*) is found by a damped Newton iteration on *F* from a self-consistent estimate of the node equilibria. The maximum residual ∥*F* ∥_*∞*_ over every accepted point of every reported curve is below 10^*−*10^.

See Algorithm 1 for pseudocode of this procedure. For the full procedure, see jr wb.continuation in the code (https://uc.figshare.com/articles/dataset/Code_Archive_for_A_Whole-Mouse-Brain_Model_of_Coupled_Neural_Masses_Analysis_of_Network_Dynamics/34845648?file=70832292).

#### Why continuation rather than simulation

Due to fundamental limitations in floating-point arithmetic, unstable equilibria are effectively invisible to computers when simulating the system normally. Without enforcing equilibrium conditions to find these states, starting the system infinitesimally close to (but not exactly on) one would still diverge to some other state when running the system forward in time. The use of floating-point arithmetic implies that our calculations cannot get infinitesimally close to or exactly equal nearly the whole set of real numbers, so the chances of a random system state landing on and staying on an unstable equilibrium are effectively zero. In general, we can think of unstable equilibria as helping to organize the system dynamics, while not being directly reachable states. Solving the system to find equilibria (as shown in Equation 8) makes the system slightly easier to compute (we have half as many unknowns, and the delays drop out) and it lets us track how equilibrium states change as we vary a parameter of interest, whether or not the equilibria are stable.

Tracing the equilibria as a parameter varies is a way to find divisions where the behavior of the system is suddenly much different; this is the heart of bifurcation analysis. However, simply tracking the state variable values at equilibria as we vary a parameter does not provide all of the information we require. To see whether an equilibrium state is stable or unstable, we look to the eigenvalues of the characteristic matrix describing the system (Section 2.4). The eigenvalues indicate the dimension of the stable and unstable manifolds at an equilibrium, and their corresponding eigenvectors indicate the direction. When all eigenvalues have a negative real part, the unstable manifold has dimension zero and the equilibrium is stable. If any eigenvalues have positive real part, the equilibrium is unstable and simulations started nearby would diverge.

### 2.4 Network Stability via Beyn’s Contour Integral Method

In general, DDE systems have an infinite number of eigenvalues, but only a finite number of these lie to the right of any vertical line in the complex plane (in particular, only finitely many satisfy Re *λ >* 0). As we are primarily interested in the case where the real parts of the eigenvalues are positive (the system is unstable) or they transition from positive to negative or vice versa (a bifurcation), this gives us an area of interest that contains a finite number of eigenvalues.

To count and estimate the eigenvalues in a box at each point along an equilibrium curve found via PALC, we have used the method of Beyn (2012) to find eigenvalues of the system with positive real components. This method utilizes a contour integral around a box Ω in the complex plane to count the number of roots of the characteristic matrix Δ(*ζ*) inside the box. For the network linearized at an equilibrium *Y* ^*\**^, the characteristic matrix is

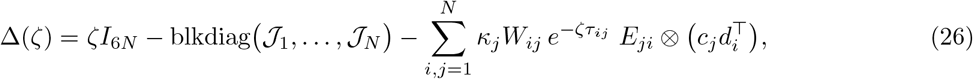

where *J*_*j*_, *c*_*j*_, *d*_*j*_ are the node-local blocks of Equation 6 evaluated at *Y* ^*\**^ and *E*_*ji*_ is the *N* ×*N* matrix with a single 1 in position (*j, i*), so that the delayed coupling from node *i* to node *j* occupies block (*j, i*). The characteristic roots *λ* are the values of *ζ* at which Δ(*ζ*) is singular; the equilibrium is stable if and only if every root satisfies Re *λ <* 0.

Beyn’s method makes use of Keldysh’s Theorem (which in turn makes use of Cauchy’s Theorem on contour integrals). Briefly, Cauchy’s theorem states that the contour integral of an analytic function around a closed loop equals zero. If the function has poles inside the loop, the integral picks up only their residues. Keldysh’s Theorem extends this to tell us what the residues of those poles are for the inverse of an analytic matrix-valued function: a pole term 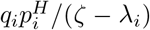 for each simple root *λ*_*i*_ (where *q*_*i*_ and *p*_*i*_ are the right and left eigenvectors of *λ*_*i*_, normalized so that 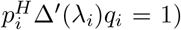 and an analytic remainder term (which contributes zero to the result due to Cauchy, and can be discarded).

We take advantage of the fact that Δ(*ζ*) is analytic (technically, entire) for DDE systems with a finite number of distinct delays and constant coefficients in Δ(*ζ*). The first criterion follows from the model construction. By linearizing the system about an equilibrium state, we achieve the second criterion. Therefore, by Keldysh’s theorem, Δ(*ζ*)^*−*1^ has poles at the roots of Δ(*ζ*), and we can find them by taking a closed contour integral. The process by which the poles are determined (Beyn’s method) involves two contour integral moments

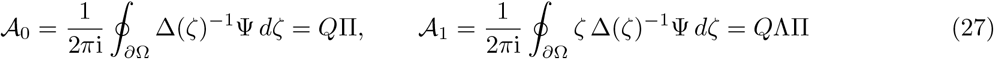

where 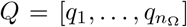 is the 6*N* × *n*_Ω_ matrix of right eigenvectors for the *n*_Ω_ roots inside the box, Λ = diag 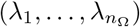, Ψ ∈ ℂ^6*N ×m*^ is a matrix of *m* random probe columns, and Π = *P*^*H*^ Ψ is the *n*_Ω_ × *m* matrix that represents how strongly each probe vector excites each root inside the box (as seen through the left eigenvectors 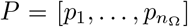. In practice we estimate these contour integrals by taking a finite sum over Gauss-Legendre quadrature nodes selected along the contour *∂*Ω.

Taking the SVD of A_0_ (truncated to rank *n*_Ω_), we can obtain Λ from A_1_ = *Q*ΛΠ by the following:

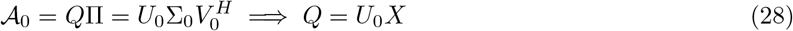

for some invertible *n*_Ω_ × *n*_Ω_ matrix *X* because *Q* and *U*_0_ span the same space. Then

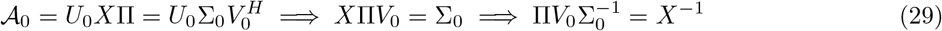

Then we see that the matrix B defined as

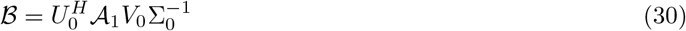

is, in fact, similar to Λ (so they share the same eigenvalues). Substituting A_1_ = *Q*ΛΠ above shows this:

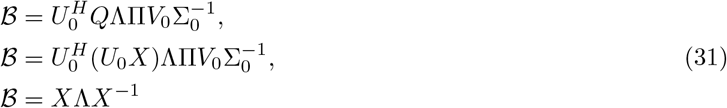

Therefore, we solve for the eigenvalues of ℬ to estimate the eigenvalues of the full network inside the contour; the columns of *X* (the eigenvectors of ℬ) give the corresponding eigenvectors *Q* = *U*_0_*X*. The eigenvalues found are only approximate because of the various approximations used above (quadrature error, truncation error of the SVD, etc.). As a result, it is necessary to polish the eigenvalues and corresponding eigenvectors with Newton’s method after each time through this algorithm. Every linear solve with Δ(*ζ*) in the contour integrals and in the polishing step uses the Woodbury identity on the block structure of Equation 26: the node-local blocks *ζI*_6_− *J*_*j*_ are inverted independently and the coupling is reduced to an *N*× *N* system, so the 6*N*× 6*N* matrix Δ(*ζ*) is never formed. The pseudocode for this procedure is contained in Algorithm 3 (and the related Algorithms 4, 5, 7, 8). For the full procedure, see jr wb.spectrum in the code (https://uc.figshare.com/articles/dataset/Code_Archive_for_A_Whole-Mouse-Brain_Model_of_Coupled_Neural_Masses_Analysis_of_Network_Dynamics/34845648?file=70832313).

#### Tracking the rightmost roots along a branch

Running Beyn’s method at every accepted continuation point would be prohibitively expensive, so stability along a branch is determined as follows (network bifurcation analysis 1param.py in the code archive, https://uc.figshare.com/articles/dataset/Code_Archive_for_A_Whole-Mouse-Brain_Model_of_Coupled_Neural_Masses_Analysis_of_Network_Dynamics/34845648?file=70832262). At the first point of the branch, Algorithm 3 is run with the eigenvalues of the uncoupled node-local Jacobians as the seed roots, and the *n*_keep_ rightmost roots in the closed upper half-plane are retained (*n*_keep_ = 64 for the continuation in *G*; 128 for the lower branch and 256 for the upper branch of the continuation in *µ*). Between consecutive accepted points, these roots and their eigenvectors are followed by Newton’s method (Algorithm 8) along a fine path of intermediate equilibria (at least 8 substeps, each a linear interpolation of the two points corrected back onto the branch). If a root in the right half of the tracked window fails to converge, converges onto another tracked root, or crosses below the real axis, the interval is redone with twice as many substeps, up to a maximum of 1024, and if the root is still lost, the tracked set is reseeded by a fresh count with Beyn’s method. Independently of these events, a Beyn count is run every 20 accepted points and whenever the sign of the rightmost real part changes, and any root it finds to the right of the lowest tracked root that is not already tracked triggers a reseed. The Beyn box uses a left edge chosen based on a gap between the edge and the tracked real parts with clearance *δ*_gap_ = 0.05 ms^*−*1^, a quadrature tolerance *ε*_quad_ = 10^*−*6^, and accepts a polished root only if its relative residual ∥Δ(*λ*)*q*∥ _1_*/*(∥ Δ(*λ*) ∥ _1_∥ *q*∥ _1_) is below 10^*−*12^.

A point is labelled stable if the rightmost tracked root has real part below −10^*−*9^ ms^*−*1^ and unstable if above +10^*−*9^ ms^*−*1^; the number of unstable roots counts each tracked root with positive imaginary part twice (for its conjugate) and each real root once. When the tracked window does not reach into the left half-plane, this count is a lower bound, which is how the triangles in Figures 1 and 2 arise. A tracked root whose real part changes sign between two points is classified as a Hopf bifurcation if its imaginary part exceeds 10^*−*4^ rad ms^*−*1^ at both points and the change in the unstable count is fully accounted for by the crossings in that interval; otherwise it is recorded as a Hopf candidate. Real roots cross the imaginary axis only at folds and branch points, which PALC locates directly.

**Fig. 1.**
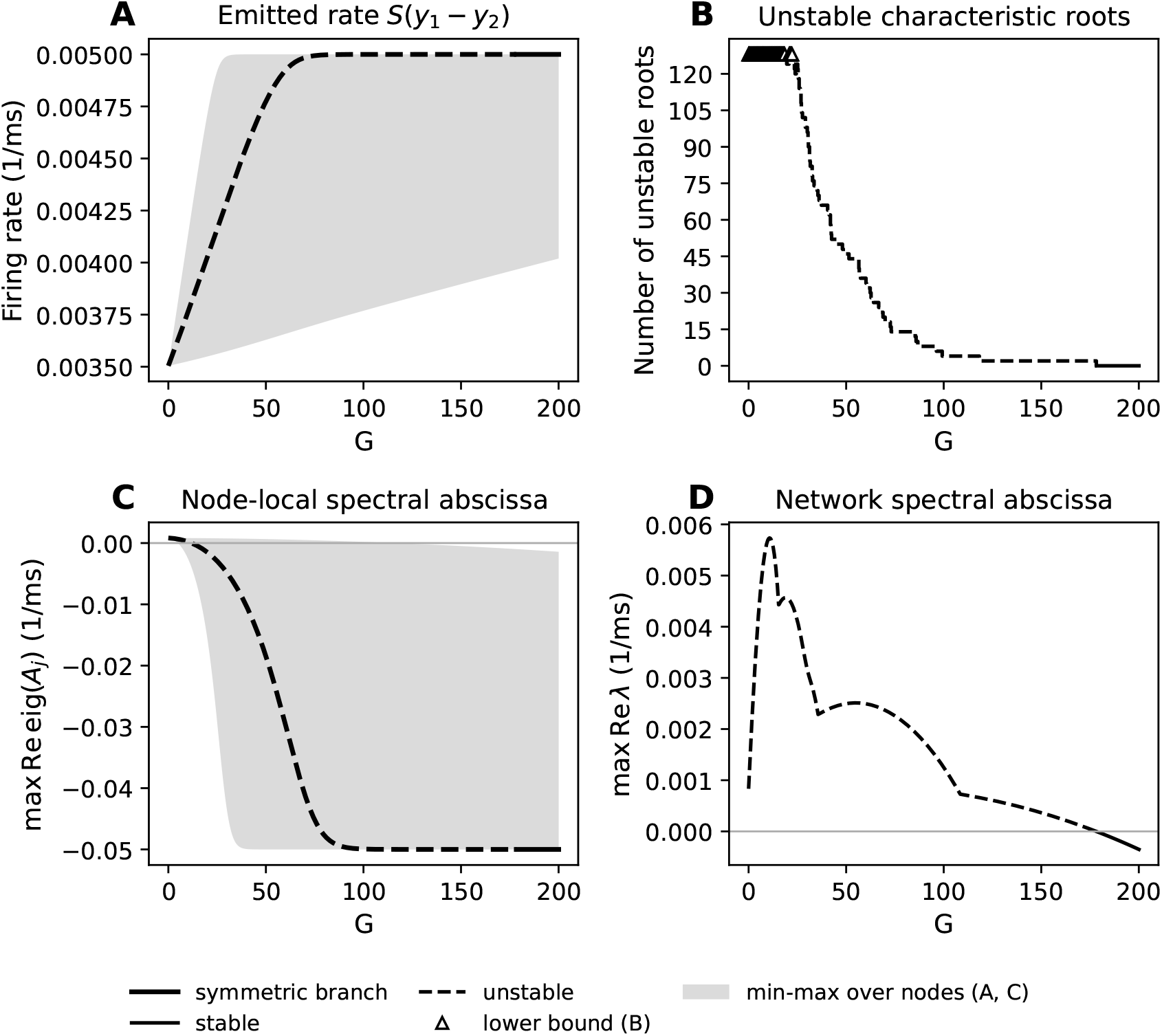
PALC in global coupling scale, *G*, at *µ* = 0.22 ms^*−*1^ and *v* = 3 m/s. In panels **A** and **C**, the median node is drawn as a black line, with line style determined by network stability: solid = stable, dashed = unstable. Also in **A** and **C**, we shade the range between the minimum value and maximum value over all nodes. **A)** Emitted firing rate *S*(*y*_1_ *− y*_2_) of the pyramidal population against *G*. The maximum value that this quantity can reach is equal to 2*ν*_max_, and this represents the sigmoid transfer function becoming saturated at high PSP values (we call a node saturated when the argument *r*(*y*_1_*− y*_2_ *−v*_0_) of its sigmoid exceeds 10). The first node saturates at *G* = 37.37, and beyond *G≈* 92 the median node is saturated. **B)** The count of unstable roots of the characteristic equation for the system. Points marked with triangles represent a lower bound on the number of unstable modes, as we only tracked 64 roots in the upper half-plane (128 counting conjugates) across the parameter range; at *G* = 0, for instance, every node contributes an unstable conjugate pair. As we can see, the relationship between *G* and the number of unstable modes is relatively straightforward. After *G ≈* 178.2, we have no remaining unstable roots and the equilibrium becomes stable. **C)** Node-local spectral abscissa, the maximum real part of all eigenvalues in individual nodes when looking only at the node-local Jacobian (see Equation 6). This effectively ignores all delays and coupling, and only looks at the instantaneous state of the system at the equilibrium point. As we can see, no individual node would continue oscillating past *G* = 116.2, while the full network remains unstable well past this point. **D)** Network spectral abscissa (maximum real part of all characteristic roots found). We see a non-monotone relationship in *G* here, with the spectral abscissa increasing from *G* = 0 to *G* = 10.71, then primarily decreasing after this point. We should note that the increases at *G* = 18.88, *G* = 54.91 and the kink at *G* = 108.3 all occur when the eigenvalue conjugate pair being tracked as the rightmost changes to a new pair of eigenvalues with a different trajectory.

**Fig. 2.**
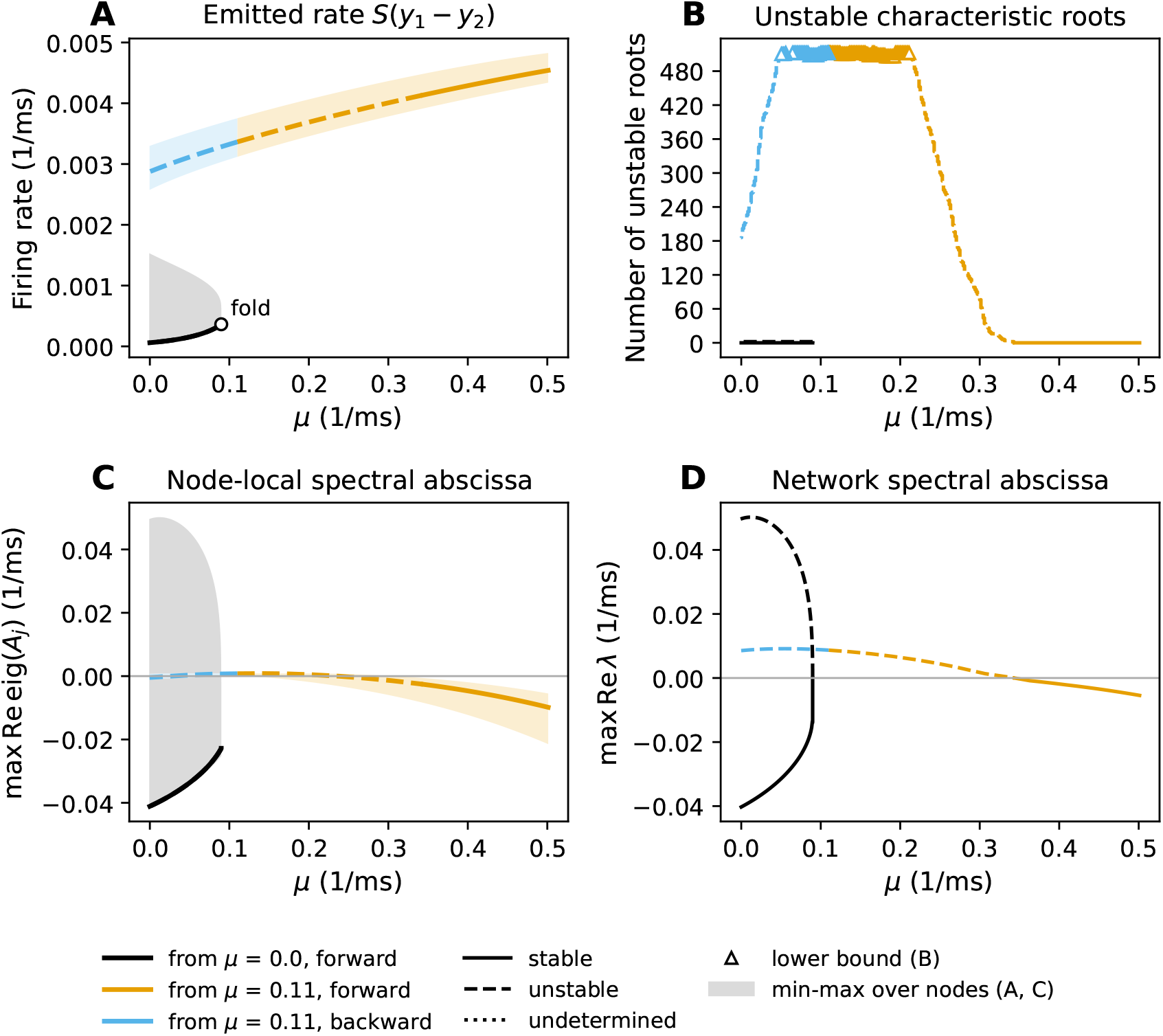
PALC in baseline external input, *µ*, for fixed global coupling scale *G* = 10 and *v* = 3 m/s. In panels **A** and **C**, the median node is drawn as a line with color determined by PALC run and line style determined by network stability: solid = stable, dashed = unstable. Three separate PALC runs were combined into this figure: starting from *µ* = 0 and continuing in the positive direction, and starting at *µ* = 0.11 and continuing in either direction. Also in **A** and **C**, we shade the range between the minimum value and maximum value over all nodes. **A)** Emitted firing rate *S*(*y*_1_ *− y*_2_) of the pyramidal population against *µ*. The maximum value that this quantity can reach is equal to 2*ν*_max_, and this represents the sigmoid transfer function becoming saturated at high PSP values. We see that starting from *µ* = 0, we found a stable equilibrium branch that continues to a fold at *µ* = 0.0898. After the fold, the equilibrium becomes unstable and continues back to *µ* = 0 (at which point we ended the continuation). Starting from a higher *µ* value, *µ* = 0.11, we find an equilibrium branch at a higher emission rate, which remains unstable down to *µ* = 0 and up to *µ≈* 0.343 (at which point the network stabilizes). **B)** The count of unstable roots of the characteristic equation for the system. Points marked with triangles represent a lower bound on the number of unstable modes, as we tracked at most 256 roots in the upper half-plane across the parameter range (512 counting conjugates; 128 roots were tracked on the low-emission branch, which never has more than two unstable roots). The low-emission branch becomes unstable at the fold with only two unstable modes, both of which are real roots (this is not a Hopf bifurcation, so the unstable state does not oscillate). **C)** Node-local spectral abscissa, the maximum real part of all eigenvalues in individual nodes when looking only at the node-local Jacobian (see Equation 6). This effectively ignores all delays and coupling, and only looks at the instantaneous state of the system at the equilibrium point. **D)** The network spectral abscissa (maximum real part of all characteristic roots found). Shows the two transitions between stable and unstable equilibria found here.

### 2.5 Maximum Lyapunov Exponent Estimation

To assess chaotic dynamics in the system, we use the method of Farmer (1982) to estimate the maximum Lyapunov exponent (MLE), *λ*_L_, of the system under various modifications. This method was chosen because it only requires solving one trajectory and a time-varying perturbation based on the local, linear dynamics of the system rather than solving two trajectories. Given the large size of the network and the computational expense of simulations, this is considerably more efficient. For a delay system the perturbation is a function on the delay interval, not a vector, so the perturbation carries its own history buffer of perturbed emitted rates, and the norm used for renormalization is taken over the current perturbed state together with that buffer (Algorithm 9).

A perturbation *δY*_*j*_ to the first-order state of node *j* (Equation 4) obeys

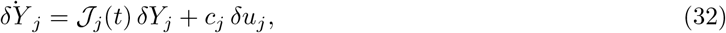

where *δu*_*j*_(*t*) = *κ*_*j*_ *Σ* _*i*_ *W*_*ij*_ *δs*_*i*_(*t* −*τ*_*ij*_), 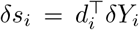, and *J*_*j*_(*t*), *c*_*j*_, *d*_*i*_ are the node-local Jacobian, input sensitivity, and emitted-rate gradient of Equation 6, evaluated along the trajectory.

Left alone, the perturbation would potentially grow/shrink exponentially and eventually overflow/underflow. As such, we renormalize *δY* and its full history every *T*_*r*_ = 100 ms. The factor by which we divide at the *k*-th renormalization, *g*_*k*_, is called the growth factor of the perturbation. After skipping an initial transient segment of the simulation to allow the system to settle onto an attractor, we track the growth factor over time. By accumulating *l*_*k*_ = ln *g*_*k*_ (for chaos, we need exponential growth) as we follow the trajectory, then dividing by the time over which we have accumulated, we arrive at the estimate *λ*_L_ of the MLE. All runs reported here use *T* = 60 s with *T*_trans_ = 20 s, so the measurement window is 40 s and contains 400 renormalizations. The model runs in ms, so *λ*_L_ is computed in ms^*−*1^; we report it in s^*−*1^.

To ensure that we have converged to an estimate, we check the following criteria for success. First, we record the log-growth at each renormalization (*l*_*k*_ = ln *g*_*k*_), and accumulate these into the cumulative log-growth 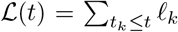 over the measurement window *t* ∈ [*T*_trans_, *T* ], so that *λ*_L_ = L(*T* )*/*(*T* − *T*_trans_). By computing the range of the residual ℒ (*t*) −*λ*_L_ (*t* −*T*_trans_) over the window, we find how consistently the trajectory has followed its final slope. We call an estimate chaotic if

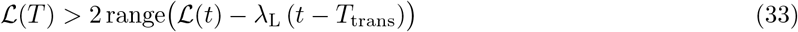

because the growth should accumulate faster than the range expands if we are converging. Similarly, we call an estimate decaying (stable) if

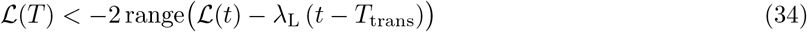

If the perturbation stays bounded (as on a limit cycle), we expect

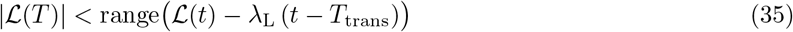

and we call the estimate periodic. Estimates between one and two times the range are reported as unresolved.

We also compute the same quantities for the first and second halves of the measurement window independently and classify each half by the same rule. A half may be unresolved while the full window is definite, since it has half the growth, but if either half is classified as chaotic or decaying and that classification disagrees with the full-window classification, the estimate has not settled and we report the realization as unresolved. The threshold of two times the range was chosen because, on a limit cycle, the log-growth is bounded however long the run is, while real exponential growth or decay accumulates linearly with the window length, so lengthening the run separates the two cases.

These tests are always performed multiple times from random initial history functions (20 realizations in every ensemble reported here; see Section 2.7) and statistical hypothesis testing (Welch’s *t* test with Bonferroni correction for multiple comparisons) is performed to detect differences between modification families in terms of MLEs. As a control, the same procedure was run on a single uncoupled node (*N* = 1): all 20 realizations were classified periodic with |*λ*_L_| *<* 0.05 s^*−*1^, as expected for the isolated node on its limit cycle.

See Algorithm 9 for pseudocode of this procedure. For more details, see jr wb.lyapunov in the code archive for the full implementation (https://uc.figshare.com/articles/dataset/Code_Archive_for_A_Whole-Mouse-Brain_Model_of_Coupled_Neural_Masses_Analysis_of_Network_Dynamics/34845648?file=70832295).

### 2.6 Synchrony and Metastability via Kuramoto Order Parameter

The Kuramoto order parameter *R*(*t*) is defined as follows:

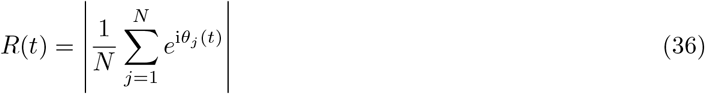

where *θ*_*j*_ is the angle of the analytic signal

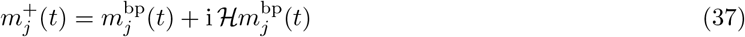

for *m*_*j*_(*t*) = *y*_1,*j*_(*t*) −*y*_2,*j*_(*t*) the pyramidal membrane potential of node *j*, 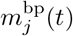 its demeaned, bandpassed version (within ± 10% of the dominant frequency), and ℋ the Hilbert transform.

*R*(*t*) is the modulus of the mean over nodes of the unit phasors 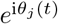. It lies between 0 (phases spread uniformly around the circle) and 1 (all nodes at the same phase), so it measures how synchronized the node phases are at time *t*. Taking the time average of *R*(*t*) gives the quantity we call synchrony here, and its standard deviation over time gives the quantity we call metastability (Deco et al., 2017).

The signal processing is as follows. Each simulation is run for *T* = 40 s at Δ*t* = 0.5 ms, the first 14 s are discarded as transient, and the remaining 26 s of *m*_*j*_(*t*) are kept at every second step (sampling rate *f*_*s*_ = 1000 Hz). The dominant frequency *f*_0_ is the peak of the Welch power spectrum (segments of 2048 samples, so the frequency resolution is 0.49 Hz) of the network-mean signal within 1–100 Hz. Each node’s demeaned signal is then band-passed with a fourth-order Butterworth filter between 0.9*f*_0_ and 1.1*f*_0_, applied forward and backward for zero phase distortion, and the phase is taken from the analytic signal. Three cycles of *f*_0_ are trimmed from each end of the phase series to remove filter and Hilbert-transform edge effects before *R*(*t*) is computed. Before any phase is extracted, a realization is checked for divergence (non-finite values) and for quiescence (median over nodes of the temporal standard deviation of *m*_*j*_ below 10^*−*3^ mV, in which case the network sits on an equilibrium and phases would be meaningless); such realizations are excluded from the synchrony statistics. No realization in the analyses reported here diverged or was quiescent.

See Algorithm 10 for pseudocode implementing this procedure. For more details, see jr wb.synchrony in the code archive (https://uc.figshare.com/articles/dataset/Code_Archive_for_A_Whole-Mouse-Brain_Model_of_Coupled_Neural_Masses_Analysis_of_Network_Dynamics/34845648?file=70832319).

### 2.7 Connectome Modification Families

The chaos (Section 2.5) and synchrony (Section 2.6) analyses ask which features of the connectome are responsible for the dynamics we observe. We answer this by comparison: the unmodified connectome is run as a *null ensemble*, and alongside it we run six *modification families*, each of which removes or randomizes one feature of the pair (*W, τ* ) while holding the others fixed. A family is a map (*W, τ* ) → (*W*^*′*^, *τ* ^*′*^) applied to the weights and delays of the network under test, with *τ*_*ij*_ = *D*_*ij*_*/v* at the default conduction speed *v* = 3 m s^*−*1^.

Writing ε= {(*i, j*) : *W*_*ij*_ ≠ 0} for the set of directed edges, every family preserves ε: no edge is created or removed, so the families differ from the null only in how weight and delay are distributed over the existing edges. The atlas is dense (|ε|= 177,018 of the 434 ×433 possible ordered pairs, or 94%), so this constraint is weak and the shuffles below redistribute over nearly every pair of regions. The six families are implemented in jr wb.connectivity modifications and are as follows.

#### Delay shuffle

The delays are permuted over the edges and the weights are left unchanged: 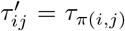 for a uniformly random permutation *π* of ε, and *W*^*′*^ = *W* . The multiset of delays is preserved exactly, but the relation between a delay and the distance it came from, and therefore the spatial organisation of the delays, is destroyed.

#### Weight shuffle

The weights are permuted over the edges and the delays are left unchanged: 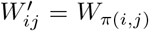 for a uniformly random permutation *π* of ε, and *τ* ^*′*^ = *τ* . The distribution of edge weights is preserved e xactly, but which pairs of regions are strongly connected is randomized, and with it each node’s in-strength Σ_*i*_ *W*_*ij*_ and out-strength Σ_*j*_ *W*_*ij*_.

#### Degree-preserving rewire

A weighted, directed version of the double-edge swap. Two edges (*i*_1_, *j*_1_) and (*i*_2_, *j*_2_) are drawn uniformly from ε, subject to *i*_1_, *i*_2_, *j*_1_, *j*_2_ being four distinct nodes and the crossed pair (*i*_1_, *j*_2_), (*i*_2_, *j*_1_) also lying in ε with positive delays. An amount *δ* is then moved from the chosen pair onto the crossed pair,

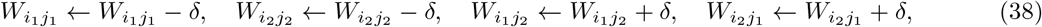

with *δ* drawn uniformly from (− min 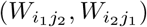, min 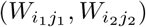) so that all four weights remain positive (a draw that would zero a weight through rounding is rejected). Each swap leaves every row sum and every column sum of *W* unchanged, so the in- and out-strength of every node, the total weight, and the support ε are preserved exactly, while the placement of weight among each node’s existing partners is randomized. The delays are unchanged. We attempt 100 |ε| swaps per realization. This is the mildest of the randomizations, since it retains all node-level statistics of the weight matrix; on the full network, the rewired matrix differs from the null by a mirror asymmetry (defined below) of about 0.33–0.35, compared with 0.88 for the weight shuffle.

#### Uniform delays

Every delay is replaced by the weight-weighted mean delay,

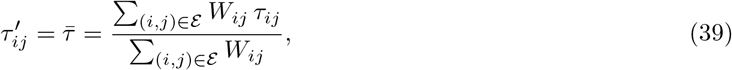

and *W*^*′*^ = *W* . The weighting is chosen so that Σ_*ij*_ *W*_*ij*_ *τ*_*ij*_ is preserved; for the full network at 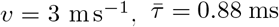 (the unweighted mean delay over edges is 1.54 ms, and the longest delay is 3.95 ms). The network becomes a single-delay system, so this family removes delay heterogeneity while retaining the delay timescale. *No delays*. Every delay is set to zero, *τ* ^*′*^ = 0, and *W*^*′*^ = *W*, so that coupling is instantaneous (in the integrator, a zero delay reads the current state at each Heun stage). Unlike the uniform-delay family, this removes the delay timescale from the network altogether.

#### Uniform weights

Every existing edge is given the mean edge weight, 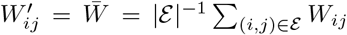 for (*i, j*) ∈ ε and 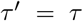 otherwise, and *τ* ^*′*^ = *τ* . The support and total weight are preserved, but the heterogeneity of edge weights, and with it the node strength distribution (and any hub structure), is removed.

Table 3 summarizes which features of (*W, τ* ) each family retains. One feature deserves comment: because the left hemisphere of *W* and *D* is the mirror image of the right (Section 2.2), the null network is symmetric under exchange of the hemispheres. The three deterministic families (uniform delays, no delays, uniform weights) preserve this symmetry, since they assign the same value to every edge of a mirror-symmetric support. The three stochastic families (delay shuffle, weight shuffle, rewire) break it. We record the asymmetry max_*ij*_ |*M*_*ij*_ − (*PMP* )_*ij*_| of *M* = *W*^*′*^ and *M* = *τ* ^*′*^, for *P* the hemisphere-exchange permutation, for every full-network realization so that this can be checked.

**Table 3.** Features of the connectome retained (✓) or removed (*×*) by each modification family. In-/Out-Degree represents the number of non-zero incoming/outgoing connection weights; “delay–distance” is the relation *τ*_*ij*_ = *D*_*ij*_ */v* between a delay and the distance it came from; “mirror” is the symmetry of (*W, τ* ) under exchange of the hemispheres.

| Family | Edge set ℰ | Weight values | In-/Out-Degree | Delay values | Delay–distance | Mirror | Random |
|---|---|---|---|---|---|---|---|
| Null (unmodified) | ✓ | ✓ | ✓ | ✓ | ✓ | ✓ | – |
| Delay shuffle | ✓ | ✓ | ✓ | ✓ | × | × | yes |
| Weight shuffle | ✓ | ✓ | × | ✓ | ✓ | × | yes |
| Rewire | ✓ | × | ✓ | ✓ | ✓ | × | yes |
| Uniform delays | ✓ | ✓ | ✓ | × | × | ✓ | no |
| No delays | ✓ | ✓ | ✓ | × | × | ✓ | no |
| Uniform weights | ✓ | × | × | ✓ | ✓ | ✓ | no |

##### Ensembles and seeds

Each family is run as an ensemble of 20 realizations at every combination of network size *N* and coupling scale *G* under test. Realization *r* uses the pseudorandom seed *s*_0_ + *r* with *s*_0_ = 42, which sets both the random initial history (each node’s state is drawn independently at a uniformly random phase on the limit cycle of the isolated node, obtained from a 6 s warm-up of a single node followed by a 4 s recording, and held constant over the delay history) and, for the stochastic families, the permutation or swap sequence that produces the modified connectivity; the deterministic families differ between realizations only in their initial history. The null ensemble is run with the same 20 seeds, so that realization *r* of every family starts from the same initial history as realization *r* of the null, and each family is a paired comparison with the null under identical initial conditions. Every realization uses *v* = 3 m/s and a step of Δ*t* = 0.5 ms. The chaos analysis uses *N* ∈{434, 80, 10} (plus the *N* = 1 control of Section 2.5) and *G*∈ {1, 5, 10}, with a run length of *T* = 60 s of which the first 20 s are discarded; the synchrony analysis uses *N* = 434 and *G*∈ {1, 5, 10, 15 }, with *T* = 40 s of which the first 14 s are discarded (Section 2.6). The same seeds are used in both analyses, so the modified connectivities and the initial histories of the synchrony ensembles are identical to those of the *N* = 434 chaos ensembles.

##### Sub-networks

To test whether an effect scales with network size, the *N* = 80 cells are run on a sub-network of the full connectome. The 80 nodes are drawn once, uniformly without replacement with seed *s*_0_, and the same node set is used for every family, coupling scale, and realization, so that the choice of nodes is not confounded with the modification. The sub-network’s weights and distances are the corresponding rows and columns of *W* and *D*, and each family is applied to the sub-network after it has been extracted, so the permutations, mean weight 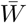, and mean delay 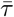 are those of the sub-network rather than of the full connectome. Because the nodes are drawn at random, the sub-network is not in general mirror-symmetric; we record the fraction of selected nodes whose contralateral homologue is also selected (25% for the *N* = 80 set and 0% for the *N* = 10 set used here).

##### Comparison with the null

For every (family, *N, G*) combination, and for every metric, the family’s ensemble is compared with the null ensemble at the same (*N, G*) by a two-sided Welch’s *t* test, with Bonferroni correction over all comparisons at a family-wise error rate of 0.05. In the chaos analysis the metric is the MLE and the correction is over the (family, *N, G*) comparisons with at least two finite estimates in each ensemble (the *N* = 1 control cells included); the comparison is reported both over every realization with a finite MLE estimate and over the resolved realizations alone. In the synchrony analysis the comparison is made for six metrics (synchrony, metastability, the mean pairwise phase-locking value overall, within, and between hemispheres, and the correlation between the simulated functional connectivity and *W* ), giving 144 comparisons for the correction, of which only synchrony and metastability are reported here (for brevity/clarity); the test uses only those realizations in which the network oscillated.

### 2.8 Computer Code

The code to run the simulations and analyses herein is written in Python (version 3.12.7) and primarily uses PyTorch (version 2.7) for computations (to enable batching on GPU). Certain analyses where GPU computation would not have made significant improvements or where PyTorch did not include necessary functions, such as statistical tests and signal processing, were run with NumPy/SciPy on CPU. All floating-point computation is in double precision. The code was initially drafted with Claude Code, utilizing the Opus 5.5 model for most tasks with periodic code reviews by the Fable 5.1 model. Following the generation of the first code draft, the authors heavily edited the code for clarity and ensured that it followed the plan exactly.

The simulations and the continuation in *µ* were run on the University of Cincinnati Advanced Research Computing Cluster (ARCC). Each simulation utilized 32 CPU cores, 416 GB RAM, and an NVIDIA A100 (40GB VRAM) GPU in Rocky Linux 8.8. To be clear, the amount of RAM was substantially more than necessary and could have been decreased to well under 100 GB (but ARCC allocates RAM based on GPU used and number of CPU cores). Equilibrium analyses did not make use of GPU, instead using Python multiprocessing on cluster nodes with 64 CPU cores; the continuation in *G* was run on a desktop workstation.

## 3 Results

The model has two parameters of primary interest when trying to match it to data: the global coupling scale *G*, which sets how strongly the connectome couples regions together, and the baseline drive *µ*, which stands in for input from unmodelled structures (thalamus, brainstem neuromodulatory systems) and is the parameter that moves a single Jansen-Rit column between rest, alpha oscillation, and seizure-like activity. Our results are organized by what can be learned about these parameters at two levels of description. The equilibrium analyses (Section 3.1) ask where in (*G, µ*) the network has a resting state at all, and whether that resting state is stable or gives way to oscillation; they are exact up to numerical tolerance, and they do not depend on initial conditions. The simulation analyses (Section 3.2) ask what the oscillatory dynamics look like once the resting state has been lost: whether they are chaotic, how synchronized the regions are, and whether that synchrony is stable or wanders in time. The first level brackets the parameter ranges worth simulating; the second tells us what a fitting procedure would be up against inside those ranges.

### 3.1 Equilibrium Analyses

A neural mass model is usually fitted to data by simulation and comparison, but simulation alone can only ever show the states the model falls into, never the structure that organizes them. An equilibrium is the network’s resting configuration: the set of regional membrane potentials at which every population’s input exactly balances its decay, with no oscillation. If that configuration is stable, the network at rest sits there, and perturbations (a stimulus, a noise burst) decay back to it. If it is unstable, the network cannot rest at it, and the equilibrium instead acts as the center around which the ongoing rhythm is organized; this is exactly the relationship between the unstable focus and the alpha limit cycle in the single Jansen-Rit column (between the Hopf bifurcations at *p* = 89.83 and *p* = 315.70, Grimbert and Faugeras (2006)). Tracking how the equilibrium moves, appears, disappears, or changes stability as *G* or *µ* varies is therefore a map of where in parameter space the model can produce quiescence, where it must oscillate, and where it jumps discontinuously between the two. Because the delays drop out of the equilibrium equations, this map can be computed at a fraction of the cost of simulation, and because it is computed by continuation rather than by forward integration, it includes the unstable equilibria that forward simulation can never reveal (Section 2.3).

See (Grimbert & Faugeras, 2006, Figure 4) for the single-node bifurcation diagram; we reproduced it with the code archive accompanying this article (single node bifurcation analysis.py, https://uc.figshare.com/articles/dataset/Code_Archive_for_A_Whole-Mouse-Brain_Model_of_Coupled_Neural_Masses_Analysis_of_Network_Dynamics/34845648?file=70832277), and the bifurcation locations quoted below agree with Grimbert and Faugeras (2006) to the precision reported there. Of primary interest to us are the two Hopf bifurcations that occur at *p* = 89.83 Hz and *p* = 315.70 Hz and the stable limit cycle that exists between them (*p* represents the average input firing rate from neurons outside of the node, in Hz). The equilibria of the node form an S-shaped curve with saddle-node (fold) bifurcations at *p* = −41.30 Hz and *p* = 113.59 Hz, so that three equilibria coexist between the folds. Between the subcritical Hopf bifurcation on the lower branch at *p* = −12.15 Hz and the Hopf bifurcation at *p* = 89.83 Hz, the node is bistable: it can rest at either a low or a high value of pyramidal PSP without oscillating, and the initial state determines which one is observed in a forward simulation. Between *p* = 89.83 Hz and *p* = 113.59 Hz the stable low-PSP equilibrium coexists with the limit cycle, and above *p* = 113.59 Hz only the oscillatory state remains. For *p >* 315.70 Hz, the system has a single stable steady state, meaning that for large values of input firing rate the system will always eventually end up on the same constant trajectory without oscillation. The limit cycle between the two supercritical Hopf bifurcations gives rise to oscillation at roughly 10 Hz.

Below we report the equilibrium curves for the full network, with continuation in *G* (with *µ* at its default of 0.22 ms^*−*1^) and in *µ* (with *G* = 10), and the stability of each curve as determined from the rightmost characteristic roots (Section 2.4), with the delays set by *v* = 3 m/s. These analyses can be thought of as direct extensions of those in Grimbert and Faugeras (2006); Touboul et al. (2011) to the many-node, delayed network case. Throughout, a node’s *effective input* means *p*_eff,*j*_ = 1000 (*µ*_*j*_ + *u*_*j*_) in Hz, the total firing-rate input to its pyramidal population at the equilibrium, which is the quantity that plays the role of *p* in the single-node diagram.

#### 3.1.1 Continuation in global coupling *G*

Here, we ask: as the connectome is turned up from zero, where does the coupling become strong enough to fully stop oscillations? This gives us a bracket on *G* indicating where simulations are possible to fit to empirical data, because the data we wish to fit will never sit at a stable steady state. The results of the analysis are shown in Figure 1. A single equilibrium branch was found, with no folds or branch points over *G*∈ [0, 200]. For *G* up to approximately 178.2 (the rightmost root crosses the imaginary axis between *G* = 178.09 and *G* = 178.33), the equilibrium has at least one unstable mode. Above this point, the equilibrium is stable at the canonical parameter values of Jansen and Rit (1995). The practical reading is that the system only becomes fully stable for extremely large *G*, which is not a useful bound on its value.

The node-level picture is more informative. At *G* = 0 every node is an unstable focus (its effective input is *p* = 220 Hz, inside the single-node alpha regime). As *G* increases, the network input raises each node’s effective input, and nodes begin to cross the single-node upper Hopf point (*p* = 315.70 Hz) between *G* = 4.8 and *G* = 5.0; by *G* = 10, 150 of the 434 nodes have crossed it and 284 remain locally unstable (unstable with the network input held fixed), and at *G* = 11.87 the effective input of the median node crosses it. The network spectral abscissa (the real part of the rightmost characteristic root of the full delayed network) rises from *G* = 0 to a maximum at *G* = 10.71 and then, with a few interruptions, decreases. Taken together, *G* ≈10–12 is where the network as a whole is driven closest to the single-node upper Hopf point while remaining most strongly unstable, and we have therefore targeted *G* = 10 as our operating point of interest in many tests.

#### 3.1.2 Continuation in baseline drive *µ*

The baseline drive *µ* is what moves a single node across the Hopf and saddle-node bifurcations of Grimbert and Faugeras (2006), and so it is the parameter through which the model expresses changes in arousal, sensory drive, or the excitation-inhibition balance of the inputs to cortex. Networking the nodes changes this picture because each region’s effective drive is now *µ* plus what it receives from the rest of the brain, which depends on the state of every other region. The question is whether the single-node structure (one branch, two Hopfs, a fold) survives in the network, or whether coupling creates additional branches on which some regions sit on the high (saturated) state while others sit on the low state. This is our most direct extension of Grimbert and Faugeras (2006) to the network case. See Figure 2 for the results of this analysis.

Unlike continuation in *G*, we found that there are multiple branches when continuing in *µ* at *G* = 10. Multiple coexisting equilibria mean that the same parameters can support qualitatively different resting configurations depending on history, which matters both for fitting (the objective becomes multimodal) and for interpretation (a change in state need not imply a change in parameters). The general shape of the bifurcation diagram is similar to that shown in Grimbert and Faugeras (2006) for the single-node case. A low-emission branch is stable from *µ* = 0 up to a fold at *µ* = 0.0898 ms^*−*1^, beyond which it turns back toward *µ* = 0 as an unstable branch with two real unstable roots (a saddle, with no oscillatory instability). A separate high-emission branch exists over the whole range examined; it is unstable from *µ* = 0 up to *µ* ≈ 0.343 ms^*−*1^, where its last pair of complex roots crosses into the left half-plane (between *µ* = 0.3421 and 0.3436), and it is stable from there to the end of the continuation at *µ* = 0.5. So for 0≤ *µ <* 0.0898 a stable resting state coexists with the oscillatory regime organized around the unstable high-emission equilibrium, and above *µ* ≈0.343 the network has a single stable steady state. The latter threshold represents considerably higher input to the nodes than in the single-node case, where the upper Hopf is at *µ* = 0.3157 (equivalent to *p* = 315.70), because the network input must be added: at *µ* = 0.343 every node’s effective input exceeds 349 Hz. Along the high-emission branch, every node becomes locally stable (stable with its network input held fixed) before *µ* = 0.309, while the full delayed network remains unstable until *µ* ≈0.343. Moving along this branch, the network undergoes a long sequence of Hopf bifurcations (Figure 2B). We believe that allowing *µ* to continue to negative values will be a useful extension here (analogous to the negative values of *p* in the single-node case); this is left to subsequent analyses because determining stability along the curve is computationally expensive, and allowing negative *µ* values increases the number of points to check substantially.

### 3.2 Network Simulation Analyses

The equilibrium analyses establish that, at the canonical parameters, the network is most strongly unstable, and closest to the single-node upper Hopf point, around *G* = 10. What equilibrium analyses cannot say is what the oscillations look like there. We therefore ran simulations to measure three properties of the oscillatory dynamics.

First, we measured whether the dynamics are chaotic: a chaotic model exhibits sensitive dependence on initial conditions, so two runs that differ by numerical round-off produce different time series, individual trajectories cannot be reproduced, and gradient-based fitting sees an objective that is rough at every scale. Second, how synchronized the regions are: empirical resting-state activity is neither fully synchronized nor independent, and a model that lives at either extreme cannot reproduce functional connectivity regardless of how it is fitted. Third, whether that synchrony is itself stable or wanders in time: metastability, in the sense of transient coordination that forms and dissolves, has been proposed as a signature of healthy cortex across fMRI, MEG, and LFP recordings (Alderson et al., 2020, 2018; Baker et al., 2014; Brinkman et al., 2022; Cabral et al., 2022; Hancock et al., 2025; La Camera et al., 2019; Mazzucato et al., 2015), and its loss has been associated with structural disconnection in traumatic brain injury and Alzheimer’s disease (Alderson et al., 2018; Hellyer, Scott, Shanahan, Sharp, & Leech, 2015).

We quantify chaos with the maximum Lyapunov exponent (Section 2.5) and synchrony/metastability with the mean and standard deviation of the Kuramoto order parameter (Section 2.6). The analyses below ask how each metric depends on coupling scale, network size, conduction speed, and specific properties of the connectivity matrix.

#### 3.2.1 Chaos and its dependence on network structure

Chaos in a two-node Jansen-Rit network was reported by Huang et al. (2011) for particular parameter choices. However, we have determined that chaos arises even at the canonical parameter values of Jansen and Rit (1995) when using an empirical structural connectome to couple the neural masses. A modeller fitting a whole-brain network needs to know whether chaos is a generic condition of the network at the default operating point, and if so, what produces it.

We suggest two explanations, each with different practical consequences. If chaos comes from the specific weights and delays of the connectome used, then it is likely a property of the brain being modelled and should be accommodated by the fitting procedure (for example by fitting distributions of statistics rather than trajectories). If instead chaos appears for any sufficiently large network of coupled nodes, regardless of how they are wired, then it is a property of the Jansen-Rit node itself under delayed coupling, and could potentially be removed by changing the model.

To distinguish between these cases, we estimated the maximum Lyapunov exponent (MLE) for the full connectome and for the six modifications of Section 2.7, at three coupling strengths (*G*∈ {1, 5, 10} ) and three network sizes (*N* ∈ {10, 80, 434 }), with 20 random initial histories each. See Figure 3 for the results of this analysis. There are several interesting trends visible in the figure.

**Fig. 3.**
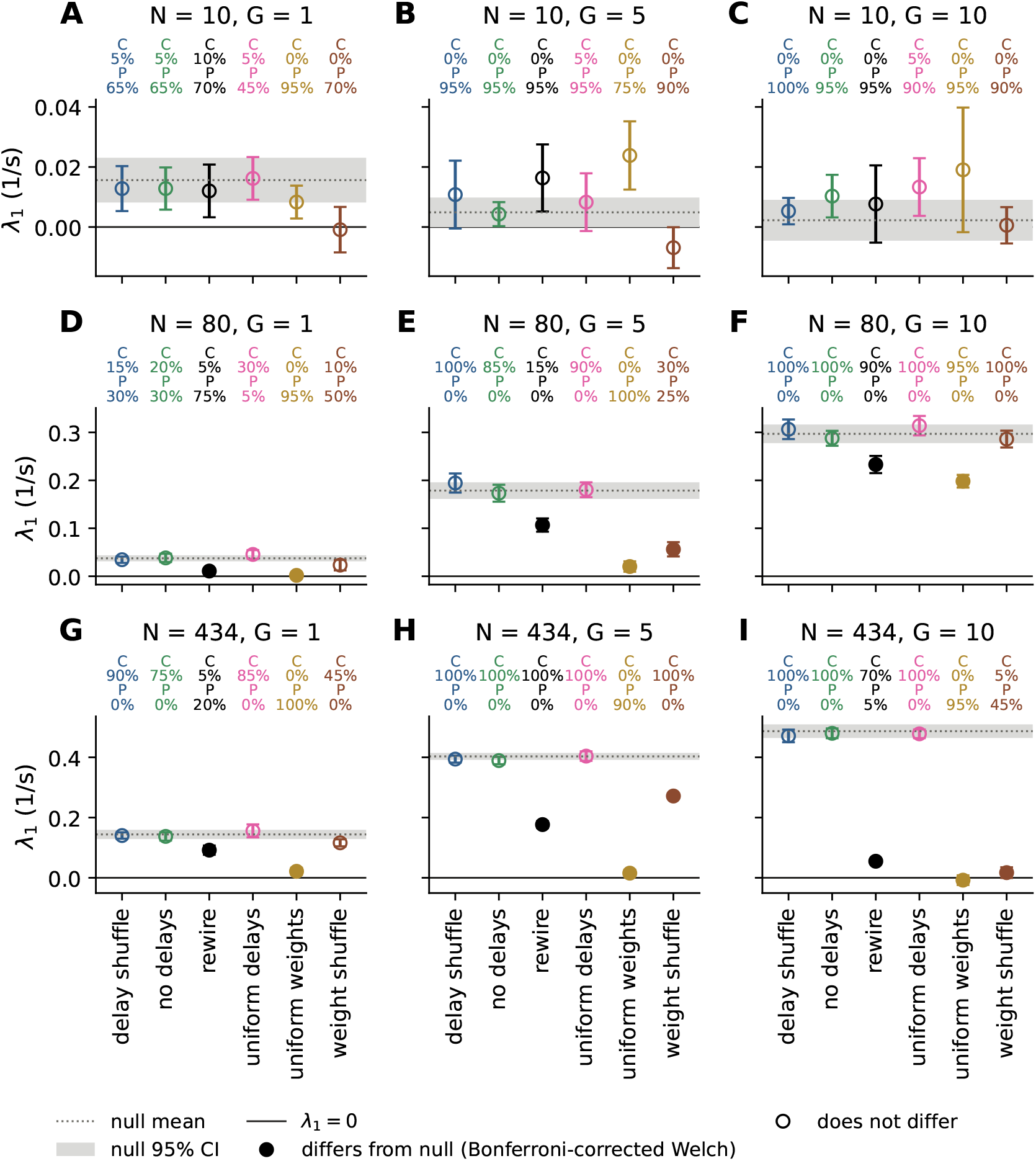
Maximum Lyapunov exponent (MLE) estimates, in s^*−*1^, by connectome modification family, coupling scale, and network size. The dotted horizontal line represents the mean value across all realizations in the null ensemble (standard connectivity matrix, as described in Section 2.2). The shaded band represents the 95% confidence interval on the mean for the null ensemble. For each connectivity modification (see Section 2.7 for details on how connectivity was modified in each case), the figure shows the mean and 95% CI on the mean across realizations, also. When a connectivity modification has changed the mean MLE estimate by an amount found to be significant (at a significance threshold of 0.05) by Welch’s *t*-test with Bonferroni correction over all comparisons, the circle showing the mean MLE is filled. Next to each connectivity modification, C indicates the percent of realizations that were classified as chaotic and P the percent classified as periodic (the remaining percent was unresolved, except for one decaying realization in panel **A**; see Section 2.5 for the classification details). Panels are grouped by number of nodes in the network in rows and global coupling strength *G* in columns. Rows of panels share a y-axis scale to make the influence of *G* more visible. There is a trend suggesting that MLE increases with both *N* and *G* for the null connectivity, and therefore in the other modifications that did not significantly impact MLE, as well. The primary takeaway is that connectivity governed by a set of uniform weights (without changing delays) is significantly less chaotic than the null for large networks, so much so that in **G, H**, and **I**, the network becomes periodic rather than chaotic in 57 of 60 realizations. The connectivity with shuffled weights (significant in **E, H**, and **I**) or rewired weights (significant in **D**–**I**), again with the same delays as the null, both cause significant decreases in MLE compared with the null for networks with many nodes, especially at strong global coupling; in **D, E**, and **G** the rewire comparison is significant only when the unresolved realizations are included. It is an interesting result that shuffling delays, making them uniform, or doing away with them entirely made no significant difference in MLE in any case.

**Fig. 4.**
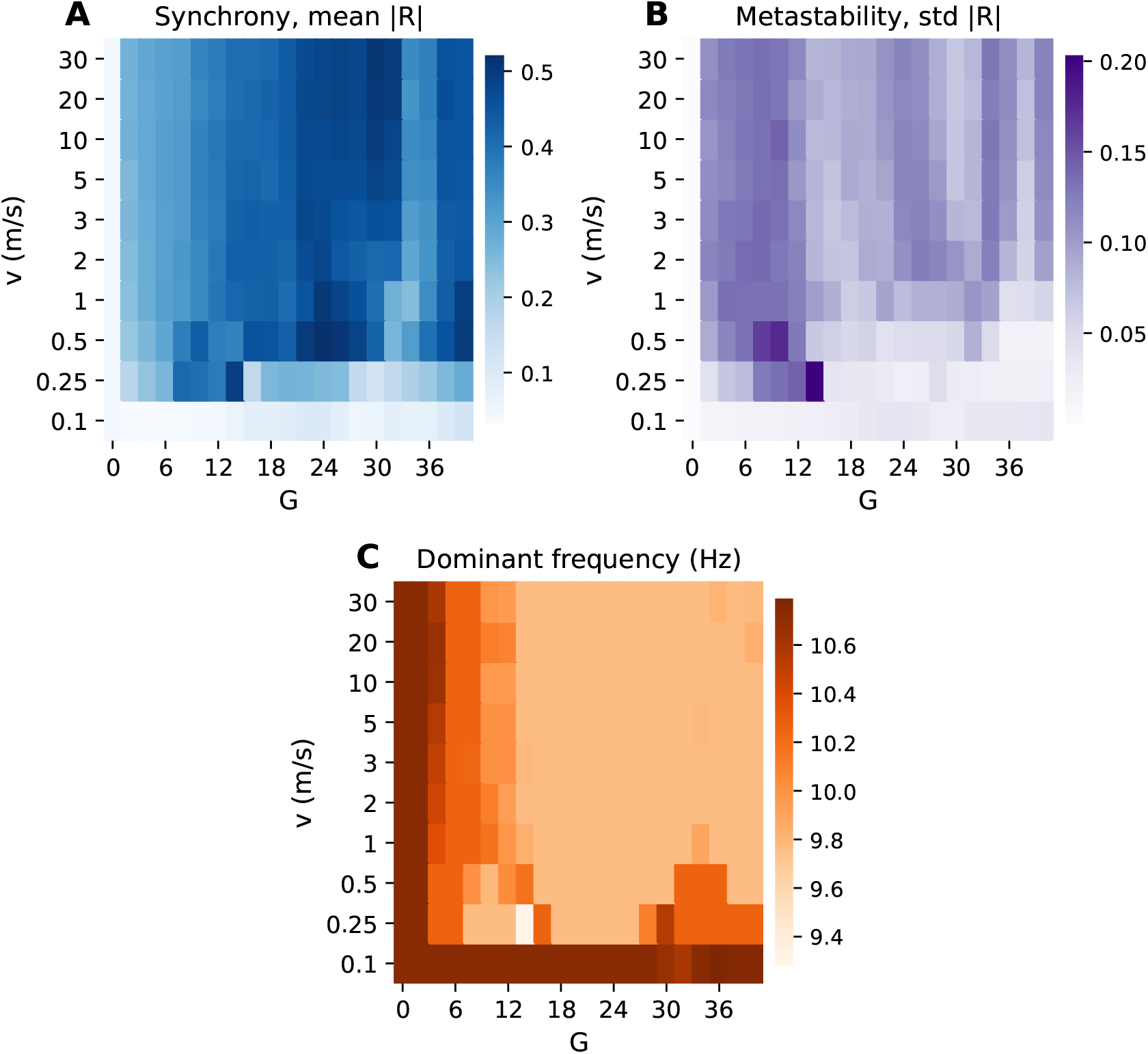
Synchrony, metastability, and dominant frequency over the (*G, v*) grid. Each cell is the mean over 20 realizations of a 40 s simulation whose first 14 s were discarded (Section 2.6); every realization oscillated. **A)** Synchrony (time mean of *R*(*t*)) increases with *G* except for a dip between approximately *G* = 30 and *G* = 36. It is hard to make any conclusion about the relationship of synchrony to conduction velocity from these data, aside from the fact that *v* = 0.1 gives near-zero synchrony (*≤* 0.12) for all *G* tested. **B)** Metastability (temporal standard deviation of *R*(*t*)) is maximal between roughly *G* = 8 and *G* = 14, especially for slow conduction velocities (*v* = 0.25 and *v* = 0.5). The peak value of metastability is 0.203 *±*0.014 (mean *±*standard deviation over realizations), and we attain this value for *G* = 14 and *v* = 0.25. The next-highest value was also for slow conduction velocity, with *G* = 10 and *v* = 0.5 giving 0.176 *±*0.018. For *G >* 14, metastability falls to about 0.03 at *v* = 0.25 and 0.05–0.06 at *v* = 0.5, and it is notable that increased *v* recovers the metastability somewhat at these high coupling strengths. **C)** The dominant frequency of the network-mean signal, averaged over realizations. The Welch estimate has a resolution of 0.49 Hz, so individual realizations take one of the values 9.28, 9.77, 10.25, 10.74, or 11.23 Hz, and the cell means lie between 9.28 and 10.79 Hz. Most grid points with *G≥* 14 sit at 9.77 Hz, and the highest cell means (10.7–10.8 Hz) occur at *G* = 0 and *G* = 2 for every *v*, and at *v* = 0.1 for nearly every *G*. The lowest frequency (9.28 Hz in every realization) occurs at a single grid point, which is the same grid point with maximal metastability (*G* = 14, *v* = 0.25). The narrow frequency range aligns with our expectation that the canonical parameter values of Jansen and Rit (1995) give roughly alpha-band EEG activity.

First, the MLE of the null ensemble (unmodified connectivity) increases with both *N* and *G*: from 0.14 s^*−*1^ at *G* = 1 to 0.49 s^*−*1^ at *G* = 10 for the full network, with every one of the 60 full-network realizations classified chaotic at *G* = 5 and *G* = 10, whereas the 10-node sub-network is periodic in nearly every realization. This suggests that increasing the number of coupled masses and increasing how strongly they communicate naturally increases the chaos in the network.

Second, the three families that alter the weights all significantly reduce the MLE in the full network at *G* = 5 and *G* = 10. The strongest effect is from the uniform-weights family, in which every edge carries the mean weight of the connectivity matrix: the full network becomes periodic rather than chaotic in 57 of 60 realizations (the remaining three are unresolved), and the effect is already significant at *G* = 1 and in the 80-node sub-network. The weight-shuffle family, which keeps the distribution of edge weights but randomizes where they sit, reduces the MLE to near zero at *G* = 10 (0.02 versus 0.49 s^*−*1^) and roughly halves it at *G* = 5. The degree-preserving rewire family also reduces the MLE substantially at *N* = 434 (to 0.18 s^*−*1^ at *G* = 5 and 0.05 s^*−*1^ at *G* = 10) and at *N* = 80. Because the rewire family preserves every node’s in-strength and out-strength exactly, the node-strength distribution alone is not what sustains the chaos; what all three families destroy is the particular arrangement of strong edges among pairs of regions in the empirical connectome. We therefore propose that this pairwise weight structure, rather than merely the breadth of node connectedness, is the primary driver of chaos in the network.

Finally, we note that the three modifications related to the delays (shuffling them, making them uniform, or removing them) had no significant impact on the MLE in any (N, G) cell. This was a rather surprising finding, as we had initially expected a combination of weight and delay structure to be responsible. In our estimation, the effects of global coupling and the structure of the weights in the connectivity matrix are considerably stronger than those related to delays, at least at the conduction speed of 3 m/s used here, where the longest delay is below 4 ms.

#### 3.2.2 Synchrony and metastability

A whole-brain model is ultimately judged on whether its regions coordinate the way real regions do. The Kuramoto order parameter gives a single number for the degree of phase coordination across the network at each instant; its mean is the synchrony, and its standard deviation over time is the metastability, the extent to which the network drifts between coordinated and uncoordinated states (Deco et al., 2017).

The two parameters that are most directly related to synchrony and metastability in a delayed network are the coupling scale *G* and the conduction speed *v*, because together they set the strength and the phase lag of every long-range interaction. We therefore mapped synchrony and metastability over a grid of (*G, v*) (Section 2.6). See Figure 4 for the results of this analysis.

Given our findings from the equilibrium continuation in *G*, we used *G* ∈ {0, 2, …, 40} because *G* = 37.37 is the point at which the first region’s sigmoid saturates at equilibrium (which is certainly a physiologically unrealistic regime; in the simulations, the saturation threshold is first exceeded at the end of the run in the *G* = 38 cells), together with *v* ∈ {0.1, 0.25, 0.5, 1, 2, 3, 5, 10, 20, 30} m/s. We expected that a value *G*∈ [10, 12] would perform best (see Section 3.1.1), because that is where the median node’s effective input reaches the single-node upper Hopf point and where the network spectral abscissa peaks; as suggested by Deco et al. (2017), neural masses close to a Hopf bifurcation perform best in terms of ability to fit empirical data. The grid search shows that our hypothesis was not far off, and an operating point near *G* = 10 or *G* = 14 is where the system exhibits peak metastability (depending on conduction velocity). We did not anticipate the peak metastability occurring at *v* = 0.25, as this is substantially lower than the default value used in The Virtual Brain and The Virtual Mouse Brain of *v* = 3 (Melozzi et al., 2017; Sanz Leon et al., 2013). We note, though, that estimates of conduction velocities in mouse vary widely between types of neurons (∼0.4 m/s up to ∼30 m/s) (Yoshida & Matsuda, 1979), complicating the selection of a single value for modeling.

The relative invariance of the measures under varying conduction velocities was also unexpected. Within the tested range of conduction velocities, for a given *G*, we mostly see differences in synchrony/metastability between roughly *v* = 0.1 and *v* = 3. This makes sense in hindsight, given the small scale of the mouse brain and the roughly 10 Hz oscillations. At *v* = 3 the mean delay (weighted by connection weight) is 0.88 ms and the longest delay is 3.95 ms; above *v* = 3 the mean delay therefore stays below 1 ms, representing roughly a 3^*°*^ phase shift at 10 Hz. While we recognize that conduction velocities around *v* = 0.25 may give more interesting dynamics, we remain unconvinced that this represents a reasonable approximation of the mouse brain.

We also repeated the same connectivity modifications (see Section 2.7) that were performed for the chaos analyses. This was intended to answer the question of whether the same properties that lead to chaos in the network are required for maximal metastability. See Figure 5 for the results of this analysis. We observe a monotone trend in *G* for synchrony in the null case, as expected: increasing the coupling scale increases synchrony (from 0.21 at *G* = 1 to 0.43 at *G* = 15). The trend in *G* is not monotone for metastability, which peaks at *G* = 10 (0.134) in the null case. This conforms well with our previous findings regarding the operating point in *G*. Importantly, the same three weight modifications that significantly reduced the MLE also differ significantly from the null in both synchrony and metastability at every *G*, with one exception (the rewire family at *G* = 5, whose metastability does not differ significantly after Bonferroni error correction). All three raise synchrony far above the null (to 0.71–0.93 for uniform or shuffled weights) and reduce metastability, to below 0.002 at *G*≥ 10 for the uniform-weights and weight-shuffle families. The three delay modifications match the null on both measures at every *G*, with one marginal exception: removing the delays entirely lowers synchrony slightly at *G* = 15 (0.414 versus 0.432, significant after correction) without changing metastability. This answers the question of whether the properties of the model that create chaotic dynamics are related to metastability: the same virtual interventions that suppress chaos also suppress metastability and push the network toward full synchrony.

**Fig. 5.**
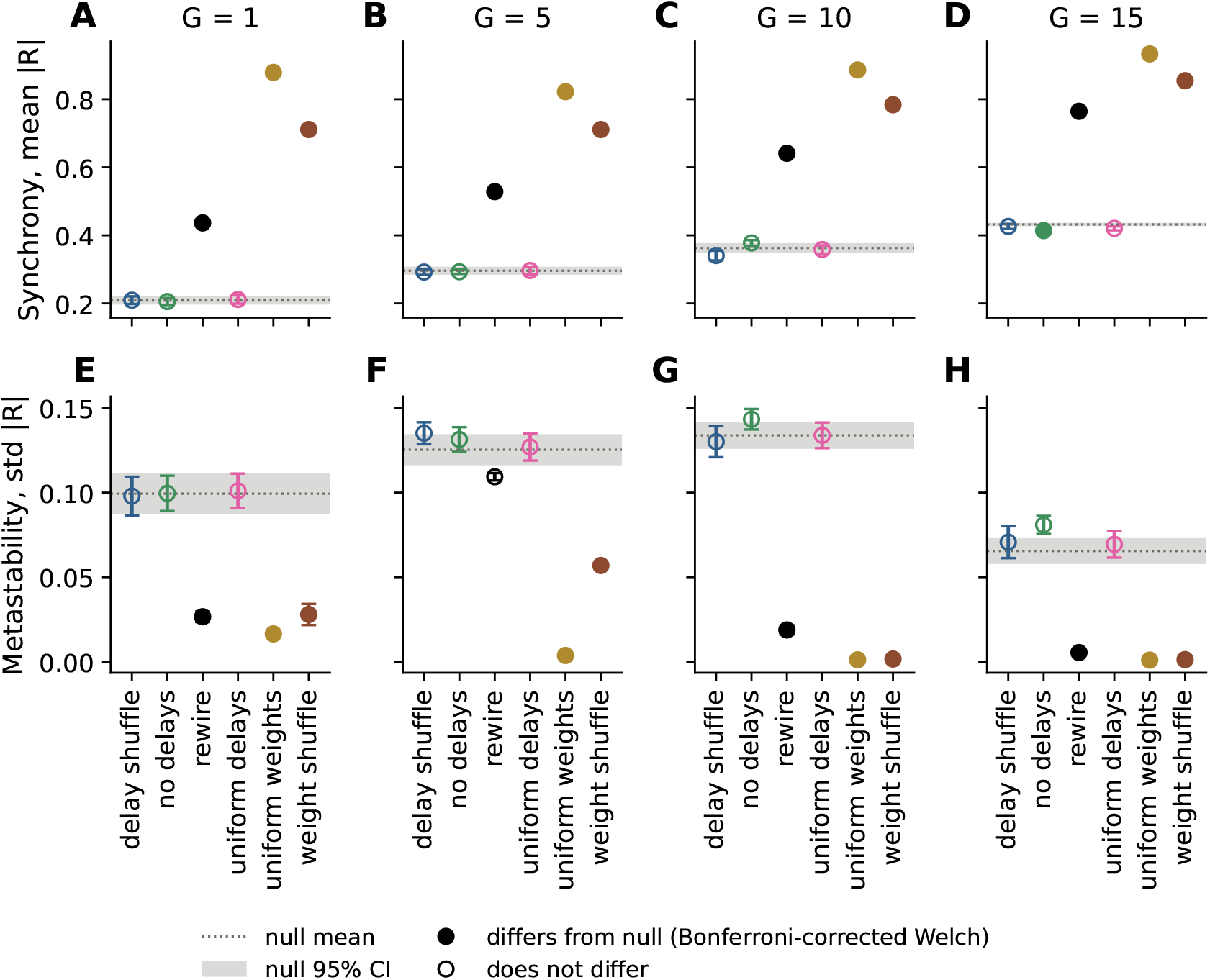
Synchrony and metastability by connectome modification family on the full network (*N* = 434, *v* = 3 m/s). The dotted horizontal line represents the mean value (synchrony or metastability, depending on the row) across all realizations in the null ensemble (standard connectivity matrix, as described in Section 2.2). The shaded band represents the 95% confidence interval on the mean for the null ensemble. For each connectivity modification (see Section 2.7 for details on how connectivity was modified in each case), the figure shows the mean and 95% CI on the mean across realizations, also. When a connectivity modification has changed the mean by an amount found to be significant by Welch’s *t*-test with Bonferroni correction over all comparisons, the circle showing mean value is filled (the Bonferroni correction includes some comparisons not reported in the figure, 144 total comparisons were used in the correction). These analyses were run with the same PRNG seeds as the *N* = 434 chaos analyses (bottom row of Figure 3, which used *G ∈* {1, 5, 10} ), which means the stochastic connectivity modifications and the initial history of each realization match the chaos analyses. **A-D)** Synchrony increases in the null case with increasing *G*, as expected. The same three weight modifications that showed significant differences from the null in Figure 3 also differ significantly here, in this case by increasing synchrony compared to the null. The uniform-weights family reached a mean synchrony of 0.93 at *G* = 15, nearly fully synchronized. Of the delay modifications, only the no-delays family at *G* = 15 differs significantly from the null, by a small reduction in synchrony. **E-H)** Metastability shows a non-monotone trend in *G* for the null case, with a peak at *G* = 10 in our simulations here. The weight modifications that significantly increased synchrony also significantly suppress metastability in every case except the rewire family at *G* = 5, which is not significant after correction; the delay modifications never differ significantly from the null.

## 4 Discussion

The problem of fitting large networks of neural mass models to empirical data, no matter what form that data takes, is a difficult one. In these analyses, we have used a structural connectome with 434 regions, which gives 434× 433 = 187,922 directed weights (177,018 of them non-zero) and as many pairwise distances (approximate tract lengths, leading to the transmission delays). Further, each of these 434 nodes has 13 parameters from the Jansen-Rit model, giving another 5,642 parameters. It is, of course, effectively impossible to perform an optimization that searches over this entire parameter space, and trade-offs must be made for the problem to become tractable. This leads us to wonder which parameters should be allowed to vary when performing the optimization, how we can best initialize the model to shorten the path the optimization algorithm must follow, and which metric(s) we should use for the optimization objective.

The analyses contained in this article represent the authors’ first attempts at answering these questions by a systematic examination of the network dynamics. The first conclusion we have drawn is that fitting based on comparing full trajectories with a metric such as mean squared error will not work, due to the chaotic dynamics. Given two runs from initial histories that differ by an arbitrarily small amount, we will obtain simulation results that are similar in frequency/amplitude of oscillations but which have completely different phases distributed among the nodes. This does not mean that the simulations are invalid, just that we need to treat model outputs in a more abstracted, statistical manner. This finding is additional confirmation of a prevailing trend in the literature: whole-brain modeling studies often fit measures such as the functional connectivity (Coronel-Oliveros et al., 2026; Wischnewski, Eickhoff, Jirsa, & Popovych, 2022), dynamic functional connectivity (Hansen, Battaglia, Spiegler, Deco, & Jirsa, 2015), or power spectral density (Coronel-Oliveros et al., 2026) between simulation and recording.

Our finding that chaotic dynamics arise, at least in part, due to the nature of the structural connectivity suggests that chaos may be a feature of the brain being modeled. This idea has been discussed extensively in the literature (Elbert et al., 1994; Faure & Korn, 2001; Korn & Faure, 2003; Rabinovich, Varona, Selverston, & Abarbanel, 2006), with many authors suggesting that chaos or transitions to/from chaos are important features of the brain. Given that alterations to the arrangement of the structural connectivity weights, and not to the delays, suppressed the chaotic dynamics in our tests, we believe that these dynamics are a fundamental property of large networks of neural masses coupled with empirical structural connectivity. The weight structure that matters is finer than the distribution of node connectedness: the degree-preserving rewire, which keeps every node’s in- and out-degree and only moves weight among a node’s existing partners, suppressed chaos nearly as effectively as shuffling or flattening the weights. The specific pattern of which region pairs are strongly connected in the atlas therefore appears to be what sustains the chaos (see Section 2.2 and Table 2 for the connectivity and some descriptive statistics).

While the authors of Hancock et al. (2025) show that metastability cannot be conclusively determined through Kuramoto order parameter methods, they acknowledge that these measures can be a useful heuristic for examining system behavior when making relative comparisons (such as between parameter regimes). Analyses of the synchrony and metastability across connectivity modifications add further evidence that the arrangement of weights in the structural connectivity matrix is vital to realistic dynamics. By removing this structure, we suppress metastability significantly and increase synchrony to near-maximal levels (Figure 5). Therefore, when fitting the model to empirical data, we cannot make simplifications that remove the chaotic dynamics in order to make the optimization problem better behaved. Doing so will, in general, also remove the network dynamics that make the model a reasonable approximation of a brain.

The impact of conduction velocity on model dynamics is rather small in the physiological range, at least on the scale of the mouse brain. At 3 m/s the weight-weighted mean delay in the network is already below 1 ms. As mentioned in Section 3.2.2, this represents a small fraction of the full period of the oscillations, and generally will cause little in the way of interference between de-synchronized nodes. This means that we believe 3 m/s is a reasonable upper bound to consider, and if the physiological value is greater than this, conduction delays must play a relatively small role in network dynamics when compared to connection strengths. The chaos analysis points the same way: at 3 m/s, shuffling, flattening, or removing the delays left the maximum Lyapunov exponent unchanged. At the other extreme, *v* = 0.1 m/s is below any viable value, as the network synchrony and metastability collapse at this speed.

The authors plan to continue investigations with this model, including extending the equilibrium analyses to find codimension-2 bifurcations. In particular, Touboul et al. (2011) showed that the single-node Jansen-Rit model exhibits a subcritical Bogdanov-Takens bifurcation as a curve of saddle-node bifurcations transitions to a curve of subcritical Hopf bifurcations (and a family of unstable limit cycles), and a curve of saddle homoclinic bifurcations. Given that period doubling homoclinic bifurcations are a common route to chaotic dynamics, we are interested in the codimension-2 bifurcations of the full network, especially in the region of the single-node Bogdanov-Takens bifurcation. We also intend to examine the behavior of the network in other configurations, including: grouping nodes based on general function and allowing each group its own coupling scale, varying certain parameters (*A/a* and *B/b*, the steady-state synaptic gains that govern the excitation/inhibition balance) globally or in functional groups, and extensions of the Jansen-Rit model such as those proposed in Wendling, Bartolomei, Bellanger, and Chauvel (2002) and Coronel-Oliveros et al. (2024, 2026).

Performing dynamical systems analyses on such a large network of delay-coupled oscillators is not only computationally expensive, there are also an extremely wide range of tests that must be run to form a comprehensive picture of the system. While we have only scratched the surface of possible analyses in this work, we believe that we have highlighted some interesting features of the system. We have determined that the global coupling scale, for our Allen Institute based connectivity, is best placed in the range *G* ∈ [10, 14] when considering metastability and the network spectral abscissa. Further, at least in the mouse brain, delays have little effect on network dynamics at conduction velocities of 3 m/s and above. Finally, the network of Jansen-Rit neural mass models at their default operating point, when governed by empirical structural connectivity data from the mouse brain, exhibits chaotic dynamics. This chaos can be suppressed by modifications of the connectivity weights, but doing so also suppresses metastability and enforces stronger synchrony.

## Supplementary information

The code archive and results are available on FigShare: https://doi.org/10.60696/uc.34845648

## Acknowledgements

The authors acknowledge funding from the National Institute on Alcohol Abuse and Alcoholism, R00AA028265 to NT.

## Declarations

The authors declare the use of generative AI technology in the preparation of this manuscript. Claude Opus was used to generate a draft of the Abstract and drafts of all algorithm pseudocode. The authors have checked the outputs for correctness, and take responsibility for any mistakes.

## Appendix A Algorithm Pseudocode

### Algorithm 1

Pseudo-arclength continuation (PALC) of equilibria *F* (*x, α*) = 0, *x* ∈ ℝ^3*N*^

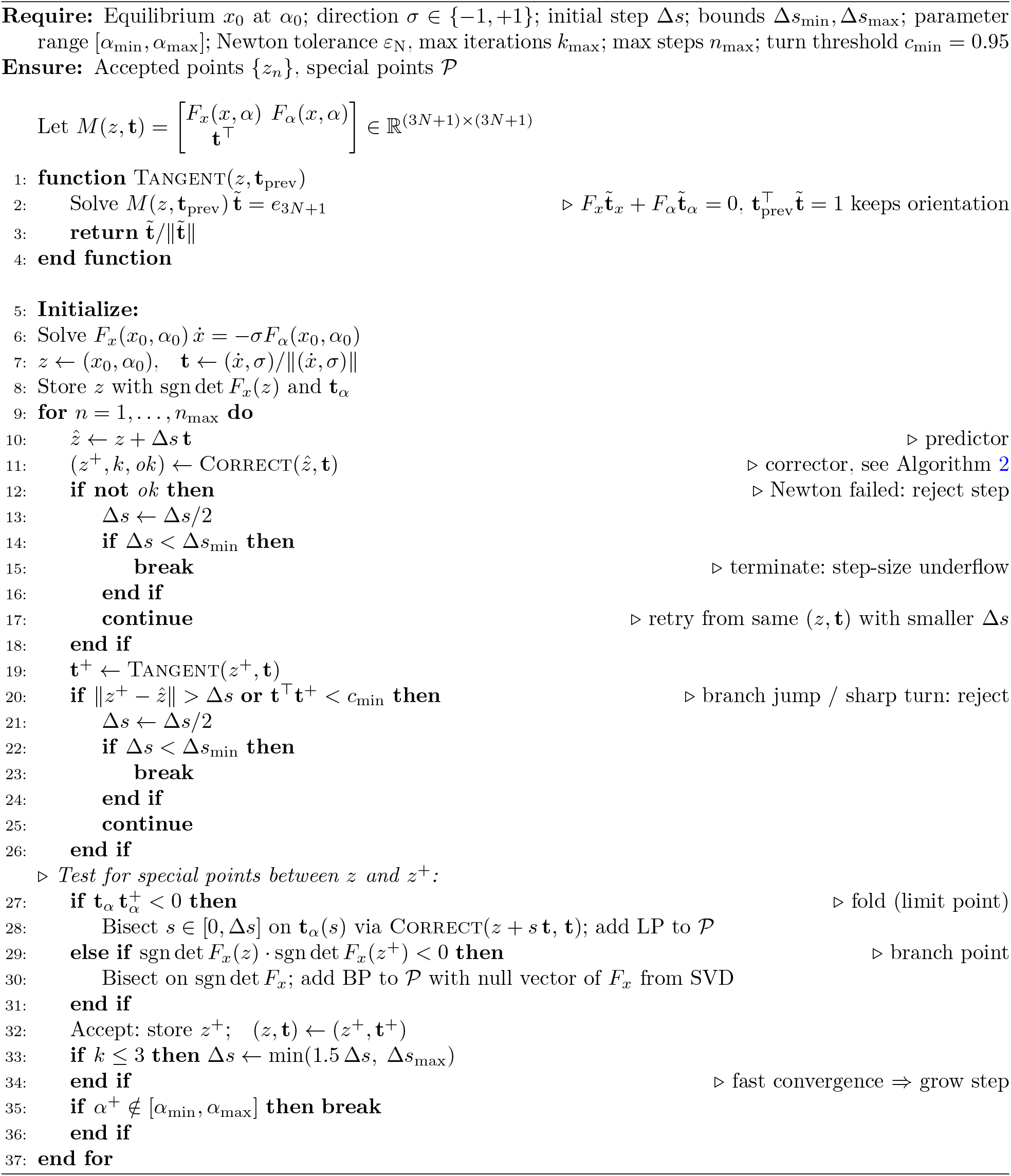

### Algorithm 2

Correct: Correct equilibrium found by PALC back onto the true curve via Newton’s method

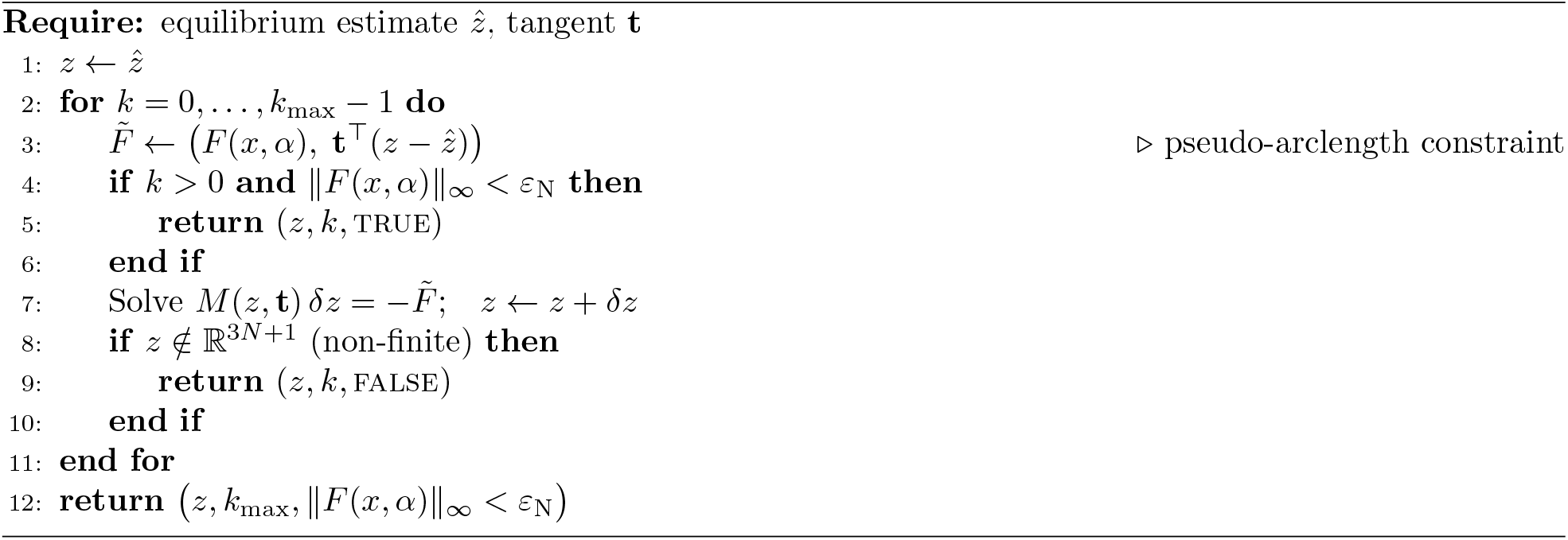

### Algorithm 3

BeynRightmost: rightmost characteristic roots at an equilibrium

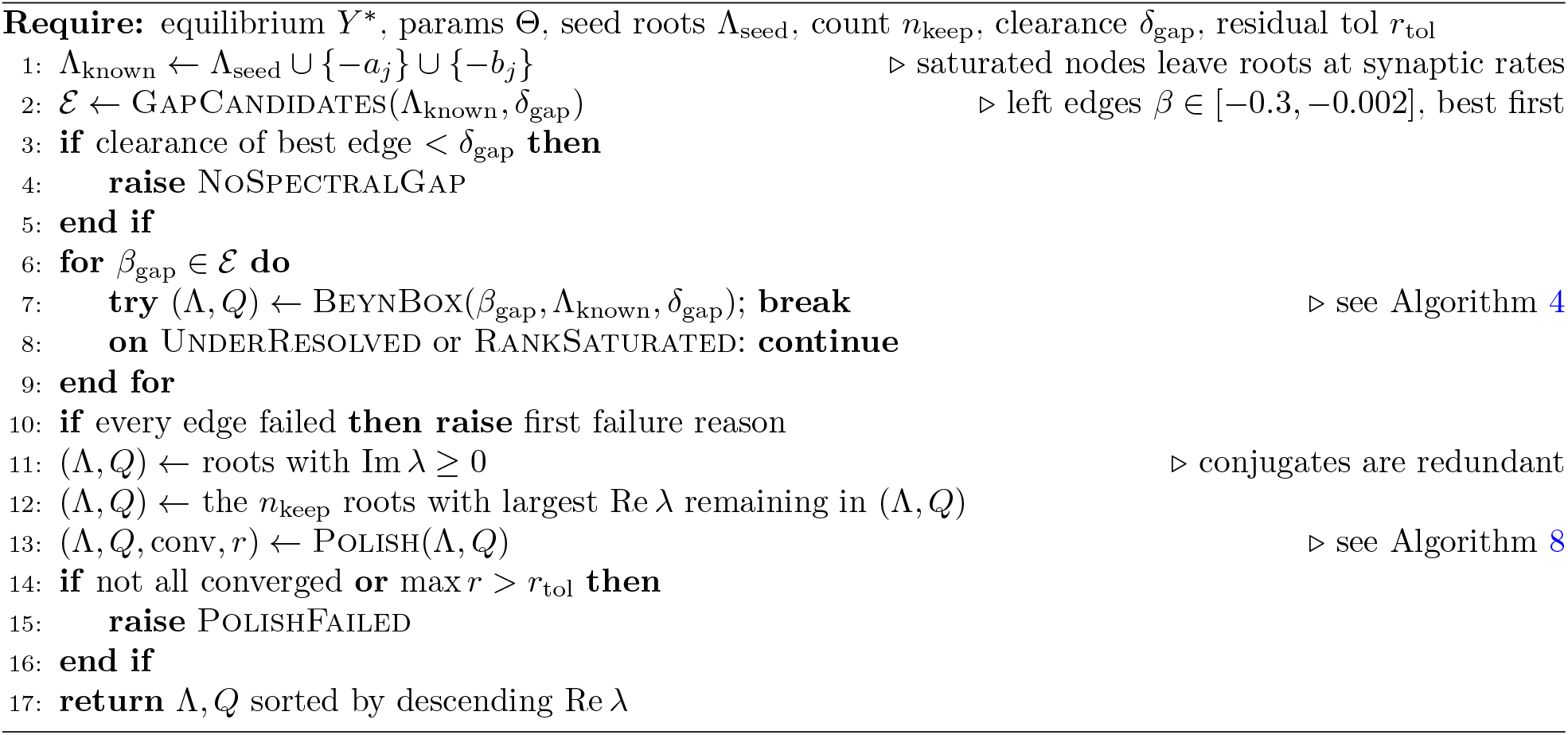

### Algorithm 4

BeynBox: retry over node counts, edge shifts and probe counts

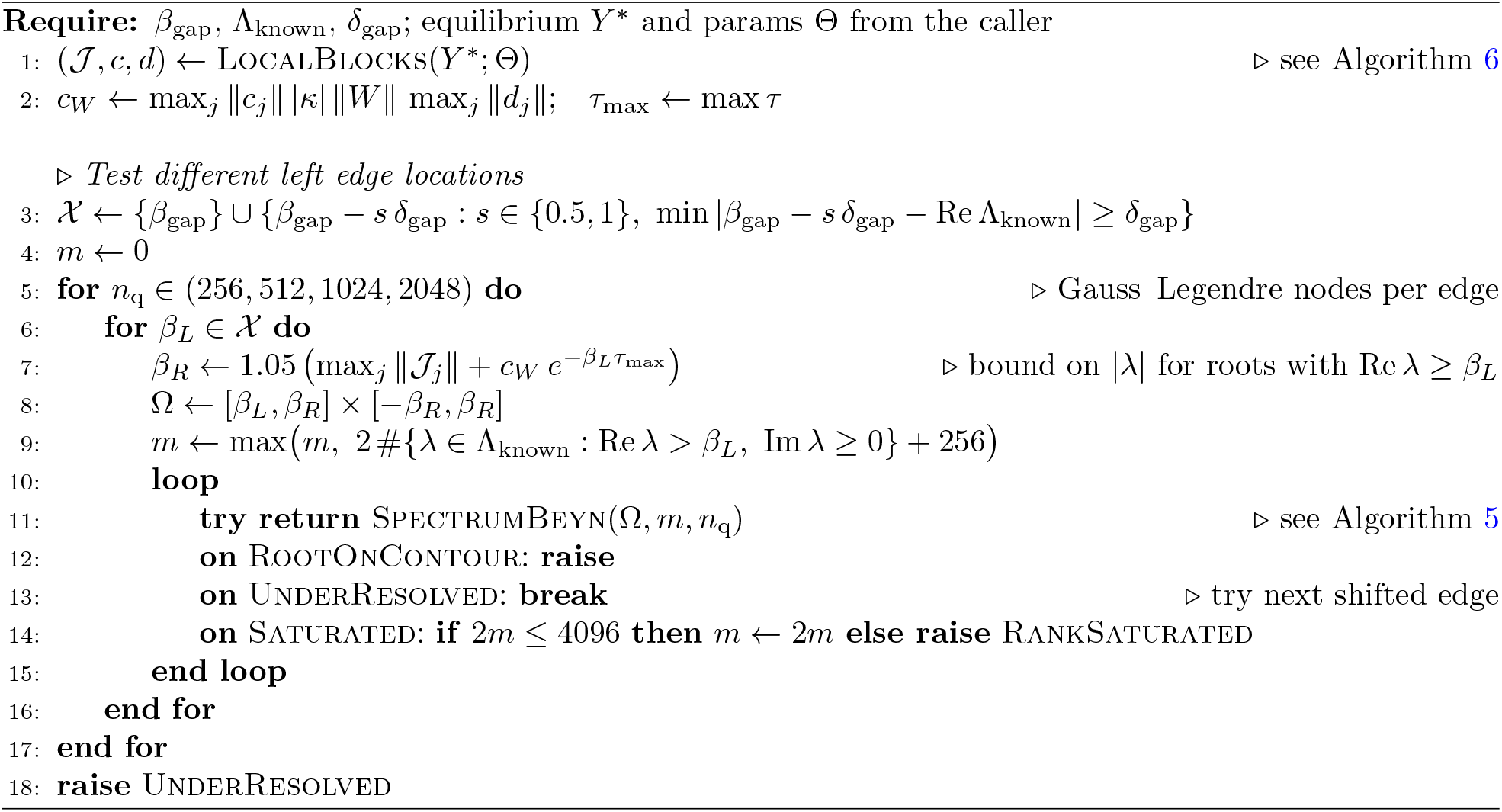

### Algorithm 5

SpectrumBeyn: Beyn’s contour-integral method on a rectangle Ω

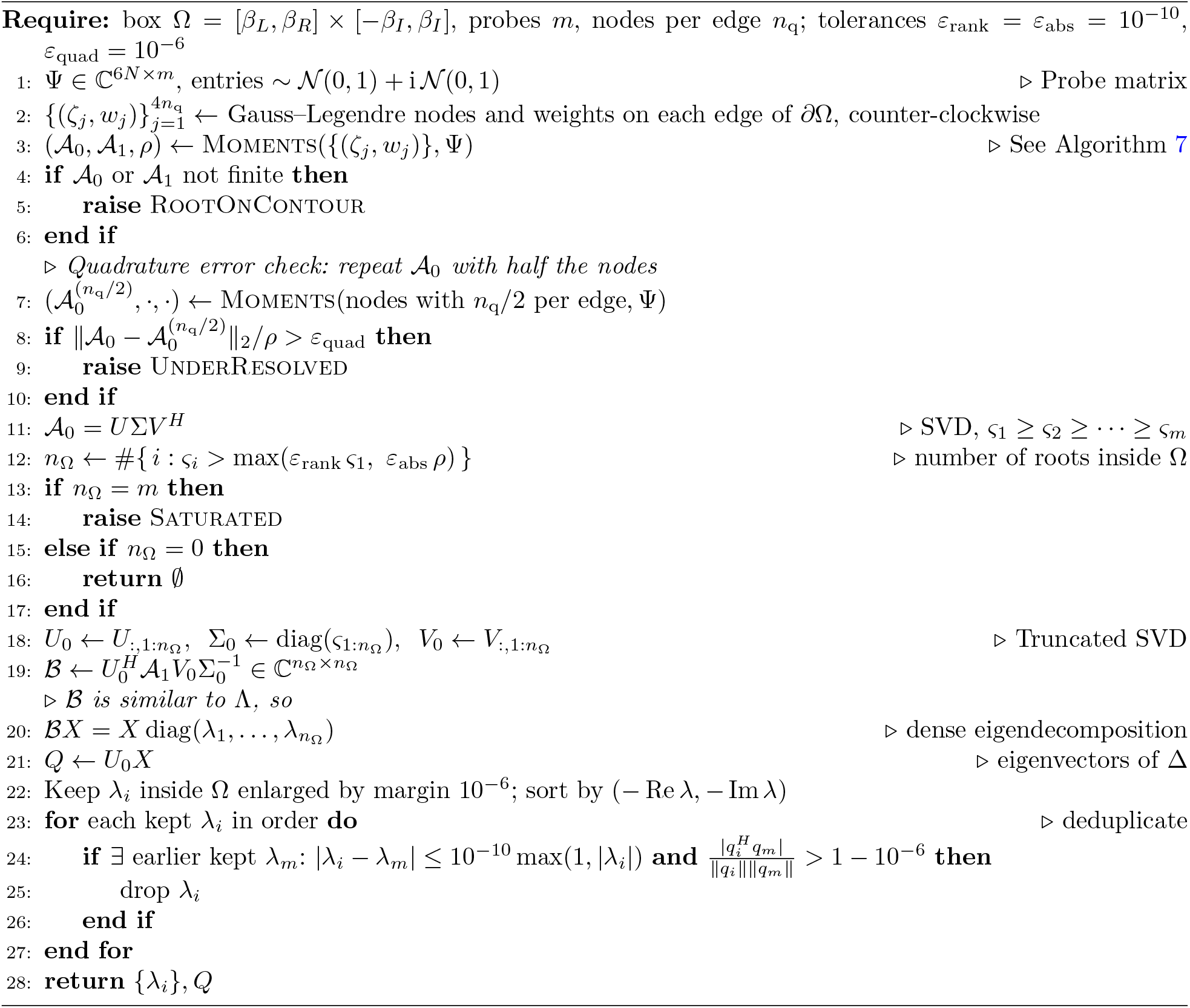

### Algorithm 6

LocalBlocks: linearize each node at state *Y*_*j*_, input *u*_*j*_ held fixed (Equation 6)

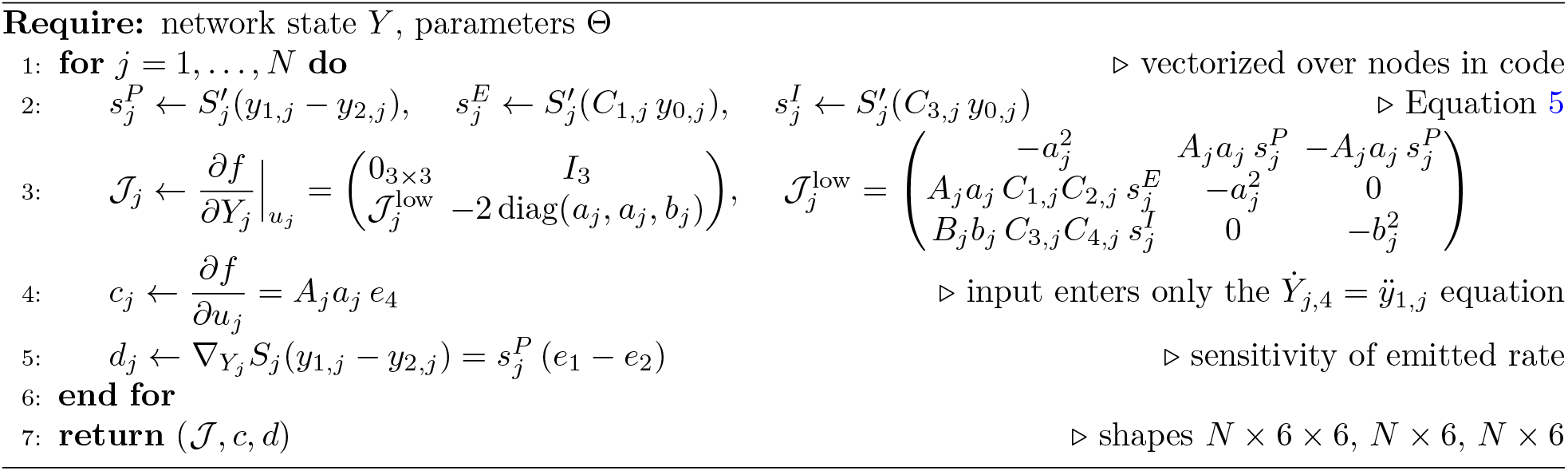

### Algorithm 7

Moments: quadrature for the contour moments

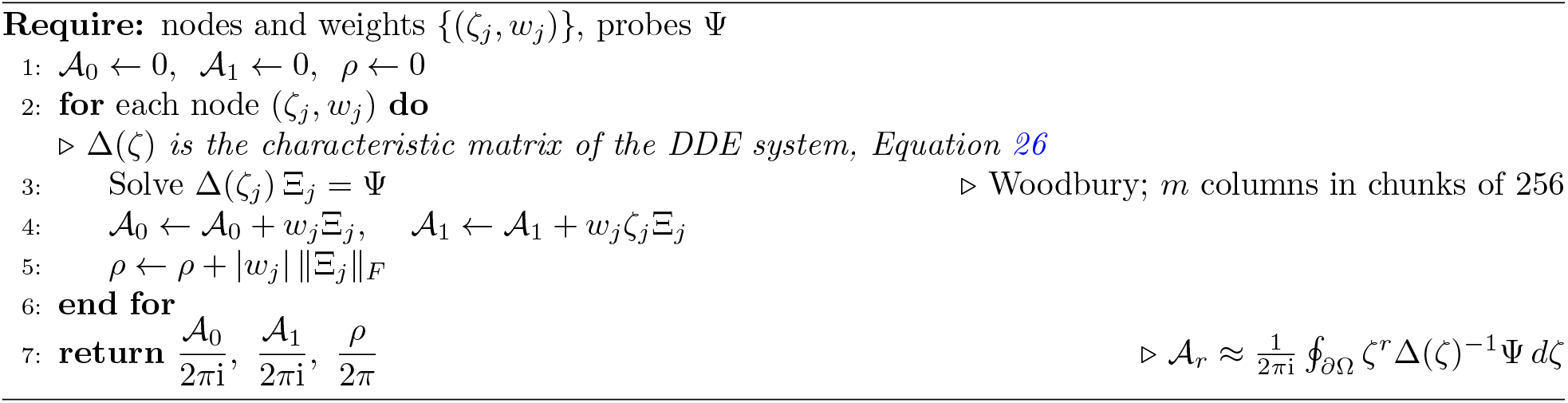

### Algorithm 8

Polish: Newton refinement of the nonlinear eigenpairs

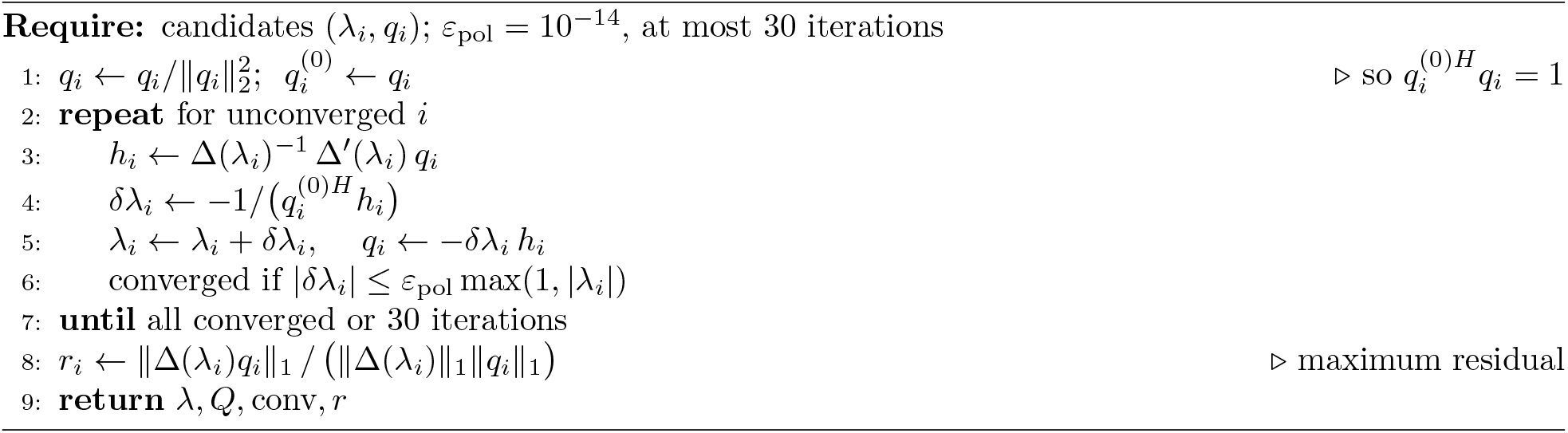

### Algorithm 9

Maximal Lyapunov exponent of the delay-coupled Jansen–Rit network (Heun integration of state + variational system, Benettin renormalization)

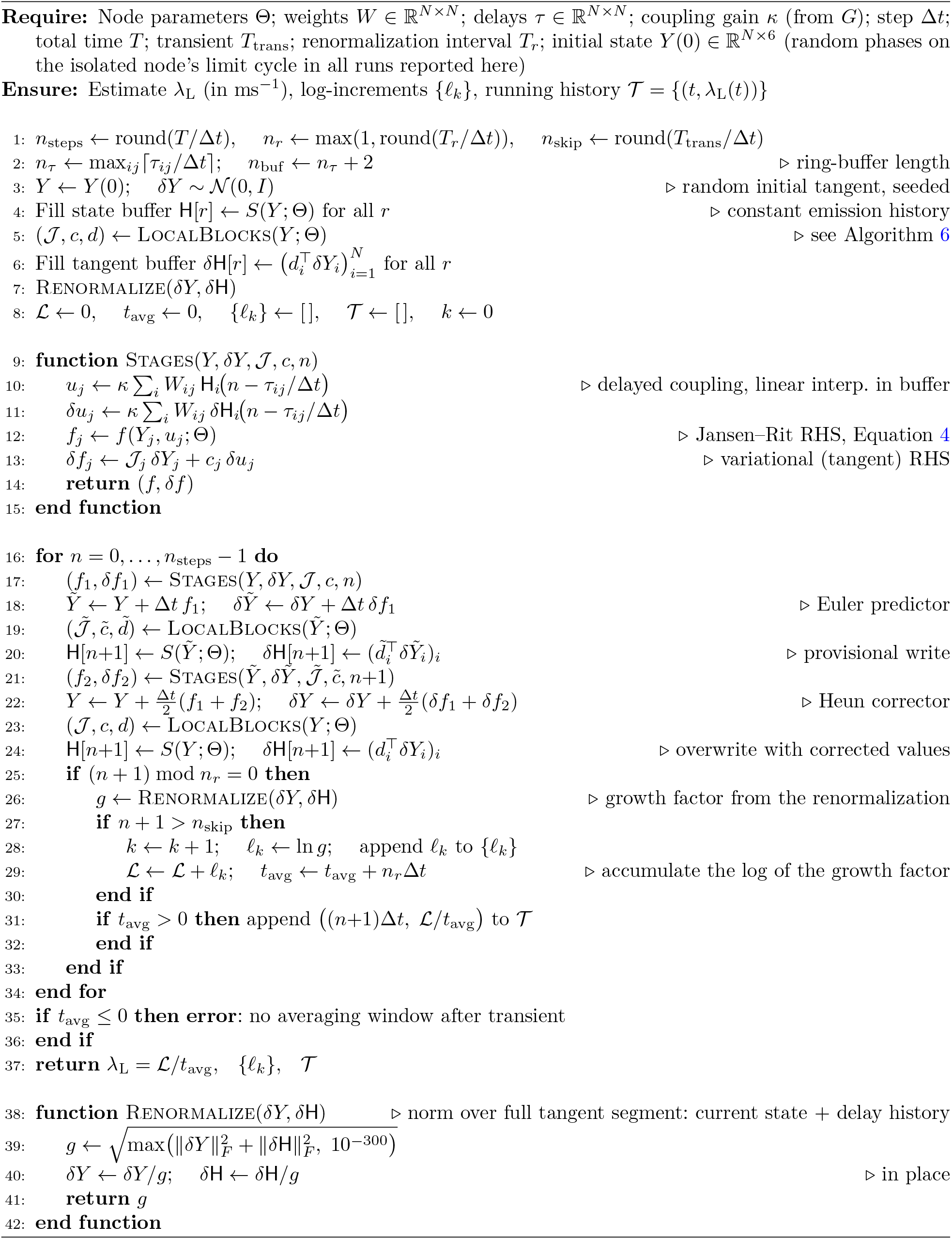

### Algorithm 10

Kuramoto R for synchrony and metastability

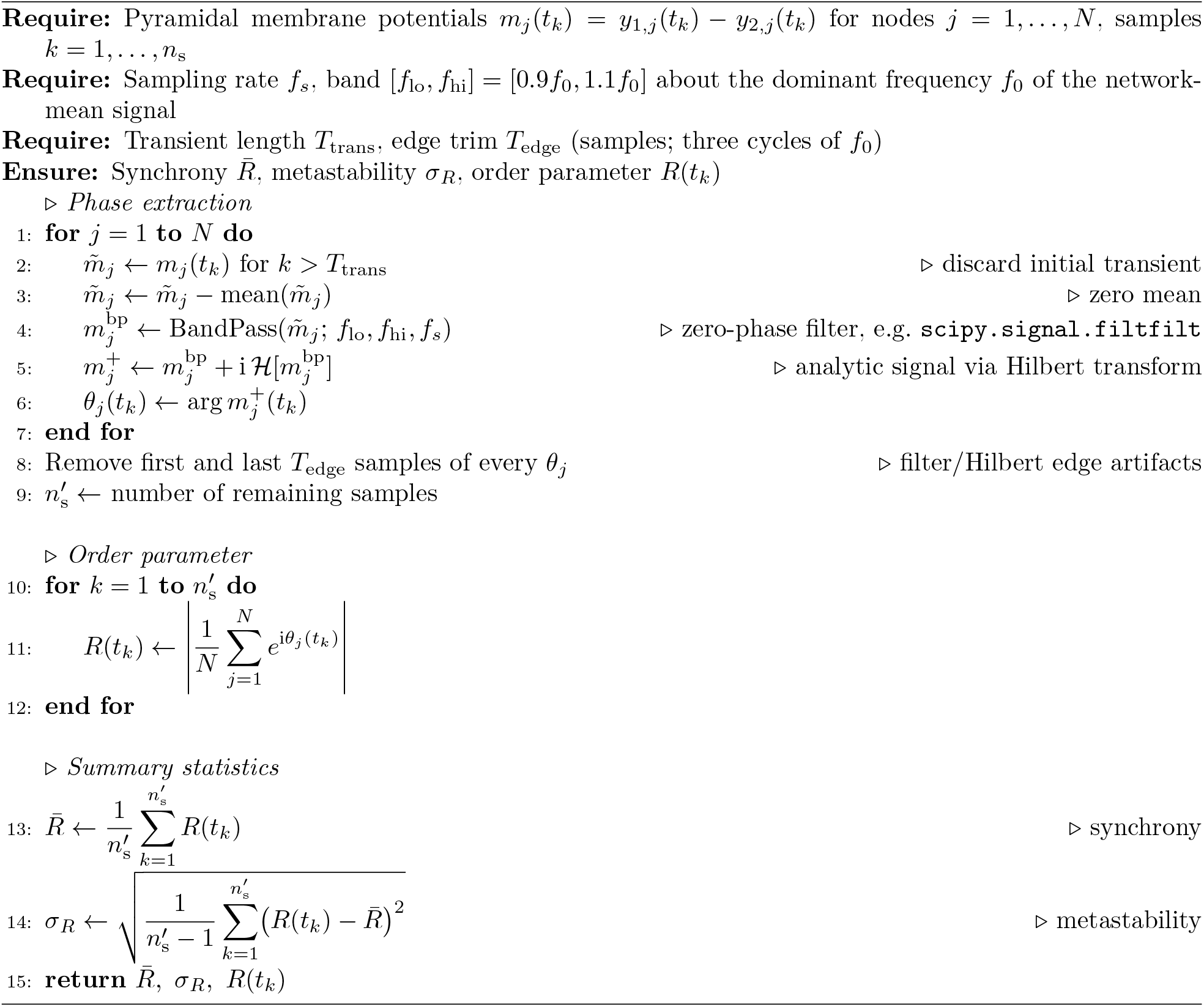

## Notes

https://doi.org/10.60696/uc.34845648

### Competing Interest Statement

The authors have declared no competing interest.

